# A *de novo* paradigm for male infertility

**DOI:** 10.1101/2021.02.27.433155

**Authors:** MS Oud, RM Smits, HE Smith, FK Mastrorosa, GS Holt, BJ Houston, PF de Vries, BKS Alobaidi, LE Batty, H Ismail, J Greenwood, H Sheth, A Mikulasova, GDN Astuti, C Gilissen, K McEleny, H Turner, J Coxhead, S Cockell, DDM Braat, K Fleischer, KWM D’Hauwers, E Schaafsma, GEMINI Consortium, L Nagirnaja, DF Conrad, C Friedrich, S Kliesch, KI Aston, A Riera-Escamilla, C Krausz, C Gonzaga-Jauregui, M Santibanez-Koref, DJ Elliott, LELM Vissers, F Tüttelmann, MK O’Bryan, L Ramos, MJ Xavier, GW van der Heijden, JA Veltman

## Abstract

**Introduction:** De novo mutations (DNMs) are known to play a prominent role in sporadic disorders with reduced fitness^1^. We hypothesize that DNMs play an important role in male infertility and explain a significant fraction of the genetic causes of this understudied disorder. To test this hypothesis, we performed trio-based exome-sequencing in a unique cohort of 185 infertile males and their unaffected parents. Following a systematic analysis, 29 of 145 rare protein altering DNMs were classified as possibly causative of the male infertility phenotype. We observed a significant enrichment of Loss-of-Function (LoF) DNMs in LoF-intolerant genes (p-value=1.00×10-5) as well as predicted pathogenic missense DNMs in missense-intolerant genes (p-value=5.01×10-4). One DNM gene identified, RBM5, is an essential regulator of male germ cell pre-mRNA splicing^2^. In a follow-up study, 5 rare pathogenic missense mutations affecting this gene were observed in a cohort of 2,279 infertile patients, with no such mutations found in a cohort of 5,784 fertile men (p-value=0.009). Our results provide the first evidence for the role of DNMs in severe male infertility and point to many new candidate genes affecting fertility.

## Main

Male infertility contributes to approximately half of all cases of infertility and affects 7% of the male population. For the majority of these men the cause remains unexplained^3^. Despite a clear role for genetic causes in male infertility, there is a distinct lack of diagnostically relevant genes and at least 40% of all cases are classified as idiopathic^3–6^. Previous studies in other conditions with reproductive lethality, such as neurodevelopmental disorders, have demonstrated an important role for *de novo* mutations (DNMs) in their etiology^1^. In line with this, recurrent *de novo* chromosomal abnormalities play an important role in male infertility. Both azoospermia Factor (AZF) deletions on the Y chromosome as well as an additional X chromosome, resulting in Klinefelter syndrome, occur *de novo.* Collectively, these *de novo* events explaining up to 25% of all cases of non-obstructive azoospermia (NOA)^3,6^. Interestingly, in 1999 a DNM in the Y-chromosomal gene USP9Y was reported in a man with azoospermia^7^. Until now, however, a systematic analysis of the role of DNMs in male infertility had not been attempted. This is partly explained by a lack of basic research in male reproductive health in general^6,8^, but also by the practical challenges of collecting parental samples for this disorder, which is typically diagnosed in adults.

In this study, we investigated the role of DNMs in 185 unexplained cases of oligozoospermia (<5 million sperm cells/ml; n=74) and azoospermia (n=111) by performing whole exome sequencing (WES) in all patients and their parents (see Supplementary Figure 1 and 2, Supplementary notes and tables for details on methods and clinical description). In total, we identified and validated 192 rare DNMs, including 145 protein altering DNMs. All *de novo* point mutations were autosomal, except for one on chromosome X, and all occurred in different genes (Supplementary Table 1). Two *de novo* copy number variations (CNVs) were also identified affecting a total of 7 genes (Supplementary Figure 3).

None of the 145-protein altering DNMs occurred in a gene already known for its involvement in autosomal dominant human male infertility. This is not unexpected as only 4 autosomal dominant genes have so far been linked to isolated male infertility in humans^5,9^. Broadly speaking, across genetic disorders, dominantly acting disease genes are usually intolerant to loss-of-function (LoF) mutations, as represented by a high pLI score^10^. The median pLI score of genes with a LoF DNM (n=17) in our cohort of male infertility cases was significantly higher than that of genes with 181 LoF DNMs identified in a cohort of 1,941 control cases from denovo-db v1.6.1^11^ (pLI male infertility=0.80, pLI controls=3.75×10^-5^, p-value=1.00×10^-5^) (Figure 1). This observation indicates that LoF DNMs likely play an important role in male infertility, similar to what is known for developmental disorders and severe intellectual disability^12,13^. As an example, a heterozygous likely pathogenic frameshift DNM was observed in the LoF intolerant gene *GREB1L* (pLI=1) of Proband_076. Homozygous *Greb1L* knock-out mice appear to be embryonic lethal, however, typical male infertility phenotypic features such as abnormal fetal testis morphology and decreased fetal testis volume are observed^14^. Interestingly, this patient has a reduced testis volume and severe oligospermia (Supplementary Notes Table 1). Nonsense and missense mutations in *GREB1L* in humans are known to cause renal agenesis^15^ (OMIM: 617805), not known to be present in our patient. Of note, all previously reported damaging mutations in *GREB1L* causing renal agenesis are either maternally inherited or occurred *de novo.* This led the authors of one of these renal agenesis studies to speculate that disruption to *GREB1L* could cause infertility in males^14^. A recent WES study involving a cohort of 285 infertile men also noted several patients presenting with pathogenic mutations in genes with an associated systemic disease where male fertility is not always assessed^16^. We also assessed the damaging effects of the two *de novo* CNVs by looking at the pLI score of the genes involved. Proband_066 presented with a large 656 kb *de novo* deletion on chromosome 11, spanning 6 genes in total. This deletion partially overlapped with a deletion reported in 2014 in a patient with cryptorchidism and NOA^17^. Two genes affected in both patients, *QSER1* and *CSTF3,* are extremely LOF-intolerant with pLI scores of 1 and 0.98, respectively. In particular, *CSTF3* is highly expressed within the testis and is known to be involved in pre-mRNA 3′ end cleavage and poylyadenylation^18^.

**Figure 1:**
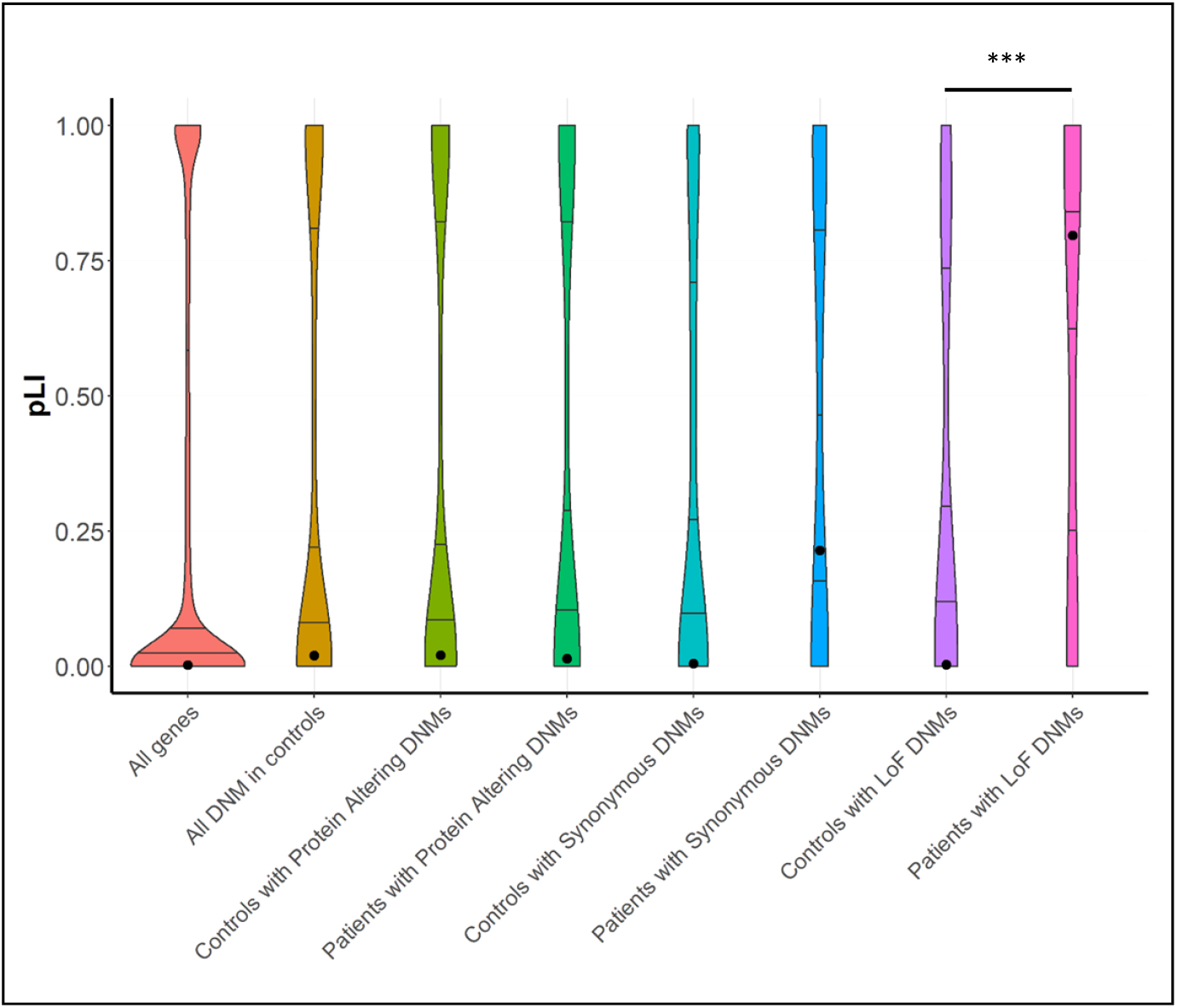
Analysis of the intolerance to loss-of-function variation for DNM genes. Violin plots represent the distribution of the pLI scores of all genes in gnomAD, all genes affected by DNMs and all LoF DNM in this study and in a control population (http://denovo-db.gs.washington.edu/denovo-db/). The observed median pLI score is displayed for each category as a black circle. The closer the pLI score is to 1, the more intolerant to LoF variation a gene is^10^. Comparison between LoF DNMs in our study and control populations shows a significance difference (p-value=1.00×10^−5^).

To systematically evaluate and predict the likelihood of these DNMs causing male infertility and identify novel candidate disease genes, we assessed the predicted pathogenicity of all DNMs using three prediction methods based on SIFT^19^, MutationTaster^20^ and PolyPhen2^21^. Using this approach, 84/145 protein altering DNM were predicted to be pathogenic, while the remaining 61 were predicted to be benign. To further analyse the impact of the variants on the genes affected, we looked at the missense Z-score of all 122 genes affected by a missense variant, which indicates the tolerance of genes to missense mutations^22^. Our data highlights a significantly higher missense Z-score in genes affected by a missense DNM predicted as pathogenic (n=63) when compared to genes affected by predicted benign (n=59) missense DNMs (p-value=5.01×10^-4^, Figure 2, Supplementary Figure 4). Furthermore, using the STRING database^23^, we found a significant enrichment of protein interactions amongst the 84 genes affected by a protein altering DNM predicted to be pathogenic (PPI enrichment p-value = 2.35 x 10^-2^, Figure 3). No such enrichment was observed for the genes highlighted as likely benign (n=61, PPI enrichment p-value=0.206) or those affected by synonymous DNMs (n=35, PPI enrichment p-value=0.992, Supplementary Figure 5). These two findings suggest that (1) the predicted pathogenic missense DNMs detected in our study affect genes sensitive to missense mutations, and (2) the proteins affected by predicted pathogenic DNMs share common biological functions.

**Figure 2:**
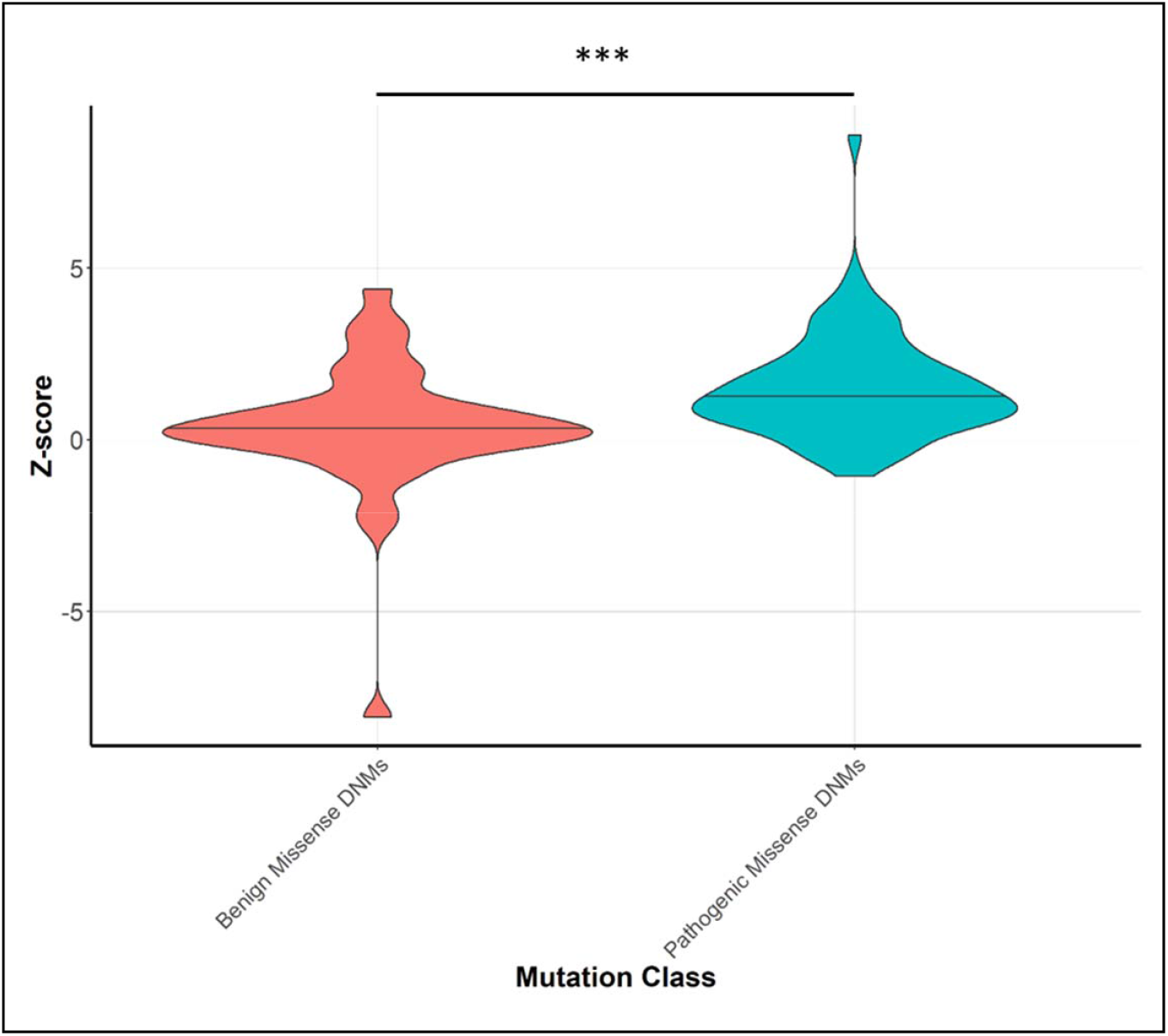
Intolerance to missense variants for genes with a DNM. Violin plots show the distribution of Z-scores of genes containing a missense DNM in our cohort, where an enrichment can be observed for predicated pathogenic DNMs in genes more intolerant to missense mutations based on their mean z-score with a p-value of 5.01×10^-4^.

**Figure 3:**
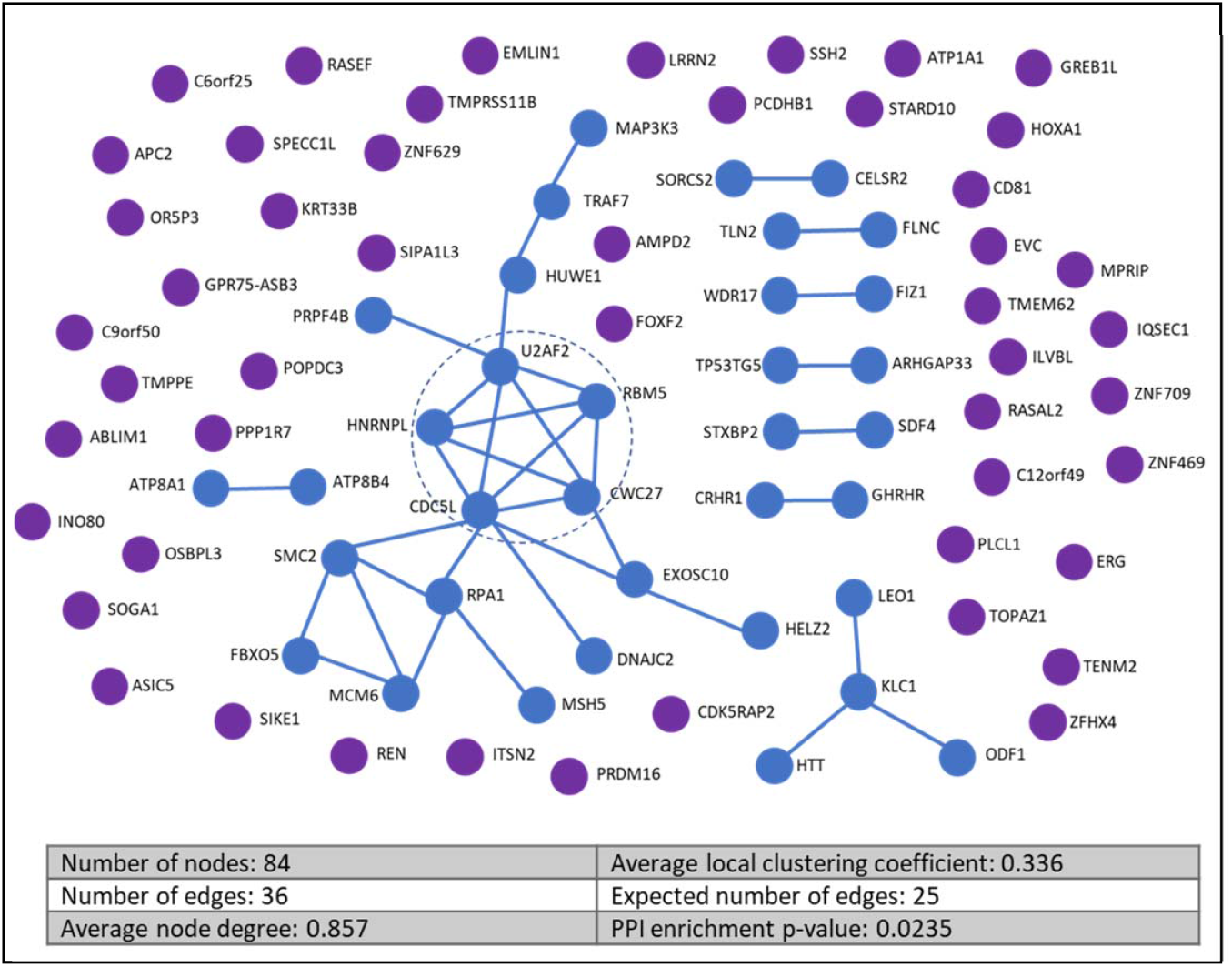
Protein-protein interactions predicted for proteins encoded by damaging DNM genes. A protein-protein interaction analysis was performed for all 84 genes containing a DNM scored as damaging using the STRING tool^23^. A significantly larger number of interactions is observed between our damaging DNM genes than is expected for a similar sized dataset of randomly selected genes (PPI enrichment p-value 2.35 x 10^-2^) with the number of expected edges being 25 and the observed being 36. The central module of the main interaction network within the figure contains 5 genes which are all involved in the process of mRNA splicing (Supplementary figure 6)

The STRING network analysis also highlighted a central module of interconnected proteins with a significant enrichment of genes required for mRNA splicing (Supplementary Figure 6). The genes *U2AF2, HNRNPL, CDC5L, CWC27* and *RBM5* all contain predicted pathogenic DNMs and likely interact at a protein level during the mRNA splicing process. Pre-mRNA splicing allows gene functions to be expanded by creating alternative splice variants of gene products and is highly elaborated within the testis^24^. One of these genes, *RBM5* has been previously highlighted as an essential regulator of haploid male germ cell pre-mRNA splicing and male fertility^2^. Mice with a homozygous ENU-induced allele point mutation in *RBM5* present with azoospermia and germ cell development arrest at round spermatids. Whilst in mice a homozygous mutation in *RBM5* is required to cause azoospermia, this may not be the case in humans as is well-documented for other genes^25^, including the recently reported male infertility gene *SYCP2^9^.* Of note, *RBM5* is a tumour suppressor in the lung^26^, with reduced expression affecting RNA splicing in patients with non-small cell lung cancer^27^. *HNRNPL* is another splicing factor affected by a possible pathogenic DNM in our study. One study implicated a role for *HNRNPL* in patients with Sertoli cell only phenotype^28^. The remaining three mRNA splicing genes have not yet been implicated in human male infertility. However, mRNA for all three is expressed at medium to high levels in human germ cells and all are widely expressed during spermatogenesis^29^. Specifically, CDC5L is a component of the PRP19-CDC5L complex that forms an integral part of the spliceosome and is required for activating pre-mRNA splicing^30^, as is CWC27^31^. U2AF2 plays a role in pre-mRNA splicing and 3′-end processing^32^. Interestingly, *CSTF3,* one of the genes affected by a *de novo* CNV in Proband_066, affects the same mRNA pathway^17^.

Whilst DNMs most often cause dominant disease, they can contribute to recessive disease, usually in combination with an inherited variant on the trans allele. This was observed in Proband_060, who carried a DNM on the paternal allele, in trans with a maternally inherited variant in Testis and Ovary Specific PAZ Domain Containing 1 *(TOPAZ1)* (Supplementary Figure 7). *TOPAZ1* is a germ-cell specific gene which is highly conserved in vertebrates^33^. Studies in mice revealed that *Topaz1* plays a crucial role in spermatocyte, but not oocyte progression through meiosis^34^. In men, *TOPAZ1* is expressed in germ cells in both sexes^29,35,36^. Analysis of the testicular biopsy of this patient revealed a germ cell arrest in early spermiogenesis (Figure 4).

**Figure 4:**
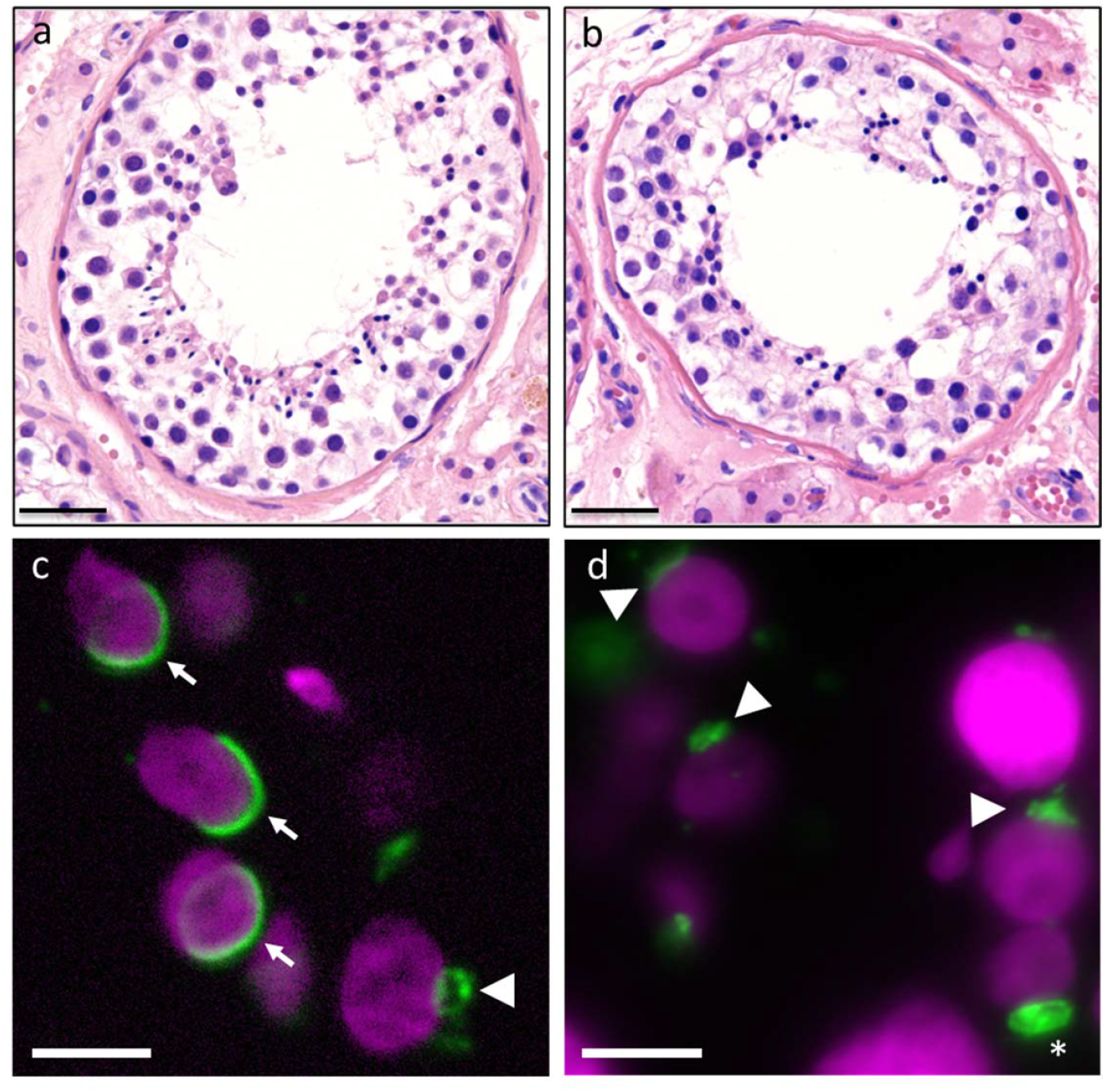
Description of control and *TOPAZ1* proband testis histology and aberrant acrosome formation: **(a,b):** H&E stainings of **(a)** control and **(b)** Proband_060 with DNM in *TOPAZ1* gene. The epithelium of the seminiferous tubules in the *TOPAZ1* proband show reduced numbers of germ cells and an absence of elongating spermatids. **(c,d):** immunofluorescent labelling of DNA (magenta) and the acrosome (green) in control sections **(c)** and *TOPAZ1* proband sections **(d)**. **(c)** The arrowhead indicates the acrosome in an early round spermatid and the arrows the acrosome in elongating spermatids. Spreading of the acrosome and nuclear elongation are hallmarks of spermatid maturation. **(d)** No acrosomal spreading (see arrowheads) or nuclear elongation is observed in the *TOPAZ1* proband. The asterisk indicates an example of progressive acrosome accumulation without spreading. Size bar in a, b: 40 μm, c, d: 5 μm.

In addition to all systematic analyses described above, we evaluated the function of all DNM genes to give each a final pathogenicity classification (Table 1, details in Material & Methods). Of all 145 DNMs, 29 affected genes linked to male reproduction and were classified as possibly causative. For replication purposes, unfortunately no other trio-based exome data are available for male infertility, although we note that a pilot study including 13 trios was recently published^37^. While this precluded a genuine replication study, we were able to study these candidate genes in exome datasets of infertile men (n=2,279), in collaboration with members of the International Male Infertility Genomics Consortium and the Geisinger Regeneron DiscovEHR collaboration^38^. The 33 candidate genes selected for this analysis include the 29 genes mentioned above and 4 additional LoF intolerant genes carrying LoF DNMs with an ‘unclear’ final pathogenicity classification. For comparison, we included an exome dataset from a cohort of 11,587 fertile men and women from Radboudumc.

**Table 1:** *De novo* mutation classification summary.

|  | Possibly causative | Unclear | Unlikely causative | Not Causative | Total |
| --- | --- | --- | --- | --- | --- |
| Missense | 21 | 38 | 50 | 13 | 122 |
| Frameshift | 4 | 8 | 1 | 0 | 13 |
| Stop gained | 1 | 3 | 0 | 0 | 4 |
| In-frame indels | 3 | 1 | 1 | 1 | 6 |
| Splice site variant | 0 | 0 | 0 | 11 | 11 |
| Synonymous | 0 | 0 | 0 | 36 | 36 |
| TOTAL | 29 | 50 | 52 | 61 | 192 |

A total of 192 rare DNMs were classified based on pathogenicity scores as well as functional data into 4 categories, 'Possibly causative', 'Unclear', 'Unlikely Causative' and 'Not causative'.

In the additional infertile cohorts, we identified only 2 LoF mutations in our DNM LoF intolerant genes (Supplementary table 2). Next, we looked for an enrichment of rare predicted pathogenic missense mutations in these cohorts (Table 2). A burden test revealed a significant enrichment in the number of such missense mutations present in infertile men compared to fertile men in the *RBM5* gene (adjusted p-value=0.009). In this gene, 5 infertile men were found to carry a distinct rare pathogenic missense mutation, in addition to the proband with a *de novo* missense mutation (Supplementary figure 8, Supplementary table 3). Importantly, no such predicted pathogenic mutations were identified in men in the fertile cohort. In line with these results, *RBM5*, already highlighted above as an essential regulator of male germ cell pre-mRNA splicing and male infertility^2^, is highly intolerant to missense mutations (missense Z-score 4.17).

**Table 2:**
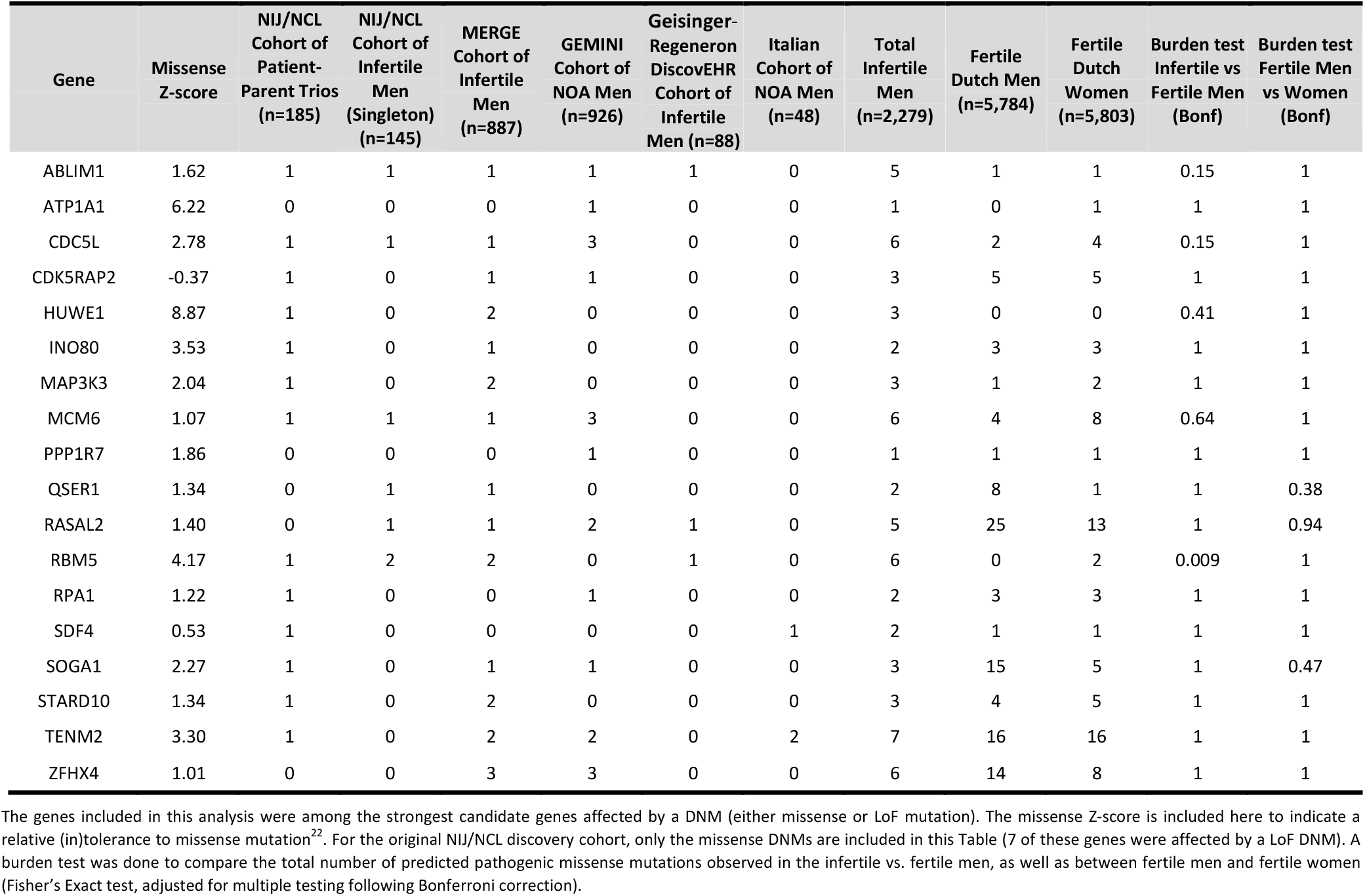
Rare potentially pathogenic missense mutations in exome data from various cohorts of infertile men and fertile control cohorts.

Given the predicted impact of these DNMs on spermatogenesis, we were interested in studying the parental origin of DNMs in our trio-cohort. We were able to phase 29% of all our DNMs using a combination of short-read WES and targeted long-read sequencing (Supplementary Table 4). In agreement with literature^39–42^, 72% of all DNMs occurred on the paternal allele. Interestingly, phasing of 8 likely causative DNMs showed that 6 of these were of paternal origin (75%). This suggests that DNMs with a deleterious effect on the future germline can escape negative selection in the paternal germline. This may be possible because the DNM occurred after the developmental window in which the gene is active, or the DNM may have affected a gene in the gamete’s genome that is critical for somatic cells supporting the (future) germline. Transmission of pathogenic DNMs may also be facilitated by the fact that from spermatogonia onwards, male germ cells form cysts and share mRNAs and proteins^43^. As such, the interconnectedness of male germ cells, which is essential for their survival^44^, could mask detrimental effects of DNMs occurring during spermatogenesis.

In 2010, we published a pilot study pointing to a *de novo* paradigm for mental retardation^45^ (now more appropriately termed developmental delay or intellectual disability). This work contributed to the widespread implementation of patient-parent WES studies in research and diagnostics for neurodevelopmental disorders^46^, accelerating disease gene identification and increasing the diagnostic yield for these disorders. The data presented here suggest that a similar benefit could be achieved from trio-based sequencing in male infertility. This will not only help to increase the diagnostic yield for men with infertility but will also enhance our fundamental biological understanding of human reproduction and natural selection.

## Data access

Raw and processed exome sequencing data of our 185 patient-parent trios is available under controlled access and requires a Data Transfer Agreement from the European Genome-Phenome Archive (EGA) repository: EGAS00001004945.

## Acknowledgements

We are grateful for the participation of all patients and their parents in this study. We thank Laurens van de Wiel (Radboudumc), Sebastian Judd-Mole (Monash University), Arron Scott and Bryan Hepworth (Newcastle University) for technical support, and Margot J Wyrwoll (University of Münster) for help with handling MERGE samples and data. This project was funded by The Netherlands Organisation for Scientific Research (918-15-667) to JAV as well as an Investigator Award in Science from the Wellcome Trust (209451) to JAV, a grant from the Catherine van Tussenbroek Foundation to MSO, a UUKi Rutherford Fund Fellowship awarded to BJH and the German Research Foundation Clinical Research Unit ‘Male Germ Cells’ (DFG, CRU326) to CF and FT. This project was also supported in part by funding from the Australian National Health and Medical Research Council (APP1120356) to MKOB, by grants from the National Institutes of Health of the United States of America (R01HD078641 to D.F.C and K.I.A, P50HD096723 to D.F.C.) and from the Biotechnology and Biological Sciences Research Council (BB/S008039/1) to DJE.

## Author contributions

This study was designed by MSO, LELMV, LR and JAV. RMS, JG, HT and GWvdH provided all clinical data and performed the TESE histology and cytology analysis under supervision of LR, DDMB, ES, KF, KDH and KM. JC performed the exome sequencing with support from BA, and bioinformatics support was provided by MJX, GA, CG and SC. Sanger sequencing was performed by PFdV, HI, HES, LEB and BKSA. MSO and HES performed the SNV analyses with support from MJX, FKM performed CNV analysis with support from AM and MSK, and GSH and LEB performed the phasing. DJE, HS, BJH and MKOB provided support on the functional interpretation of mutations. DFC, LN, CF, SK, FT, KIA, ARE, CK, and CG-J were involved in the replication study. The first draft of the manuscript was prepared by MSO, HES, RMS, MJX, GWvdH, and JAV. All authors contributed to the final manuscript.

