## Supplementary material for "A *de novo* paradigm for male infertility": Online Material and Methods

### Materials and methods

Cohort of infertile patients and fertile parent trios

We enrolled a total of 185 patients who presented with unexplained (idiopathic) azoospermia (N=111) or severe to extreme oligozoospermia (with or without asthenozoospermia N=74) at the Radboudumc outpatient clinic between July 2007 and October 2017 (N=170) and at the Newcastle upon Tyne Hospitals NHS Foundation Trust (Newcastle, UK) between January 2018 to January 2020 (n=15). The reference values and semen nomenclature were used according to the WHO guidelines[1] (see Supplemental Note). Clinical evaluation did not lead to an etiologic diagnosis and all patients were negative for AZF deletions and chromosomal anomalies (see Supplementary notes). The study protocol was approved by the respective Ethics Committees/Institutional Review Boards (Nijmegen: NL50495.091.14 version 4, Newcastle: REC Ref: 18/NE/0089, Bursa: 05.01.2015/04) and written informed consent from all patients and their parents was obtained. We used residual genomic DNA extracted from a blood sample taken at the time of evaluation and treatment at the fertility centre. DNA from all proband's parents was obtained from saliva by using the Oragene OG-500 kit (DNA Genotek, Ottawa, Canada).

Immunofluorescence staining of human testis biopsies

Tissue sections were cut from formalin fixed paraffin embedded (FFPE) testicular biopsies. As staining controls testicular biopsies obtained from fertile men after a previous vasectomy was used. FFPE sections were prepared for staining following standard protocols. To detect RBM5, antibody HPA018011 from Atlas Antibodies was used (1:500). For detection, a donkey anti-rabbit Alexa488 antibody was used (1:1000), which was applied in combination with lectin coupled to Alexa568 (1:1500) to detect the acrosome (both Thermo Scientific). Slides were counterstained with DAPI. Images were obtained with a Zeiss Axio Imager ZI fluorescence microscope equipped with the Zen software package.

Cohort of verified fertile Dutch parents

We used an anonymized exome dataset derived from 5,784 Dutch men and 5,803 Dutch women who had conceived at least one child as a control cohort for the frequency of rare variants in fertile men and fertile women. These men and women received routine exome sequencing at the Radboud diagnostics centre as the healthy parent of a child with a severe illness. Although these men fathered a child with intellectual disability, their fertility is expected to be similar to an unselected sample of the male population.

Exome sequencing

WES samples were prepared and enriched following the manufacturer's protocols of either Illumina's Nextera DNA Exome Capture kit or Twist Bioscience's Twist Human Core Exome Kit. All sequencing was performed on the NovaSeq 6000 Sequencing System (Illumina) at an average depth of 72x (Illumina's Nextera Kit) and 99x (Twist Bioscience's Kit). Sequenced reads were aligned to Human Reference Genome (GRCh37.p5/hg19) using BWA Mem v0.7.17[2], Picard[3] and GATK v4.1.4.1[4]. Following best practice recommendations, single nucleotide variations and small indels were identified and quality-filtered using GATK's HaplotypeCaller. Afterwards, all variants were further analyzed using a custom GATK4-based algorithm to identify and separate high- and low-confidence *de novo* variants from inherited variants. Briefly, posterior genotype probabilities (GQ) were recalculated for each sample at each variant site using Bayes's rule to take into account family and population priors[4,5]. Variants absent in parental samples with recalculated GQ >= 10 and allele count (AC) below 4 or allele frequency (AF) < 0.1%, whichever is more stringent were classified as low confidence DNMs. Variants with recalculated GQs >= 20 and the same AC/AF criterion were classified as high confidence DNMs. Afterwards, tagged variants with coverage < 10, variant read percentage < 15% and GATK quality scores < 400 were removed to ensure only the most reliable variants were considered. Sanger sequencing was then used to validate DNMs calls. Ensembl's Variant Effect Predictor (VEP)[6] was used to fully annotate all *de novo* variants.

Variant filtration and interpretation

The primary stages in filtering of variants included removing all variants with an allele frequency of >0.1% in the gnomAD database to only include rare variants in our analysis. All variants then with <10 reads in the exome data and/or less than 15% of these reads containing the mutation were then removed. At this stage, any remaining variants lying outside the exonic regions were then removed.  This provided the initial list of 192 rare de novo variants. All synonymous and non-protein altering spice site variants were then removed, leaving a total of 145 protein altering rare de novo mutations.

Pathogenicity prediction was then based on SIFT[7], MutationTaster[8] and PolyPhen2[9] and all variants were classified according to the American College of Medical Genetics and Genomics (ACMG) and the Association for Molecular Pathology (AMP) 2015 guidelines[10]. All protein altering variants predicted to be pathogenic by at least 2 out of 3 prediction models, absent from the fertile male cohort, present in <5 males in the gnomAD database were considered for further functional analysis (n=84).

Functional analysis was split into 6 different categories, each category provided a score of either 1 or 0 depending on whether they met the threshold for that category. These categories included: RNA expression of the gene in the testis, RNA enrichment in the testis or presence in spermatogenesis, protein expression in the testis, whether an infertile mouse model already exists for the given gene, the protein function in relation to spermatogenesis and finally whether the given gene interacts with any known fertility genes. For expression levels retrieved for each gene of interest from the GTEx database (https://www.gtexportal.org/), an expression of medium (≥10 < 100 TPM) or high (>100 TPM) gave a score of 1 with low (>2 <10 TPM) and no expression (<2 TPM) giving a score of 0. Tissue expression was retrieved from the Project The final classification of the genes was then split into 'Not causative', 'Unlikely causative', 'Unclear' and 'Possibly causative'. These classifications were given based on the variant scores out of 6 with: [0 points + "Not expressed"/"Not detected"/"Not Present" on several occasions = Unlikely causative], [0 points + "Unknown" on several occasions = Unclear], [1-2 points = Unclear] and [3-6 points = Possibly causative].

Variant validation

Validation of low-quality DNMs was performed using standard Sanger Sequencing approach to confirm the presence of the mutation in probands and its absence in the parents. Primers for each SNV were designed using PrimerZ[11] and PCR reactions were performed using AmpliTaq 360 DNA Polymerase (ThermoFisher, MA, USA) according to the manufacturer's protocol. Primer sequences and PCR conditions are available upon request.

Phasing Analysis to determine Parent-of-Origin

The origin of DNMs identified in the exomes of patients was first investigated in the short-read exome data by performing phasing analysis on those variants that contained a parental informative SNP (iSNP) within 150bp from the DNM. As a next step, all DNMs were target-enriched with long range PCR and sequenced using the Oxford Nanopore tech (ONT) MinION sequencer (Oxford Nanopore technologies, Oxford Science Park, UK). Target regions were designed to encapsulate both the DNM and a parentally informative SNP, from which parent-of-origin, and allele frequencies (percentage read counts associated to a given allele) could be ascertained and DNM pre-/post-zygosity could be ascertained.

Primers were designed using our Primer3[12] (version 2.3.6) and GRCh37.p5 based in-house GUI-wrapped pipeline. All Expected fragment sizes were limited to a maximum of 12 kb for quality control and enrichment success rate. For those DNMs with no exome supported iSNPs within a 10 kb distance, primers were designed to cover approximately 2.5 kb on either side of the DNM with the expectation of finding additional iSNPs in the intronic regions. Long-range PCR target enrichment was carried out using our optimized running conditions of 3 separate supermixes/enzymes (see supplementary table 5). Sample fragment sizes were confirmed using gel electrophoresis, and quantities were measured with the Qubit dsDNA HS kit (Thermo Fisher Scientific, Waltham, MA, USA), with the best quality supermix enrichment for each given sample/target selected for sequencing, where quality was assessed by cleanest banding in gel electrophoresis and greatest ng/ul.

The long-range PCR target enrichments of >20 ng were prepared for sequencing with the ONT ligation sequencing kit (SQK-LSK109) following the manufacturer's protocol, with adjustments for sample type and yield. Individual sample libraries were concentrated where necessary at given bead clean-up steps and pooled based on fragment size. Fragment size-based pools were combined prior to flowcell loading. Prepared samples were sequenced on the MinION using the FLO-MIN106 version 9.4 flowcell platform. Flowcells were run until complete pore exhaustion, with minimal refuel of flowcells performed whenever active pore percentages dropped below 70%, achieving an average of 30 billion basecall yields per flowcell and coverage depth per sample of >5000X.

The sequence signal data in multi-fast5 format were basecalled using Guppy[13] (version 3.4.4, https://nanoporetech.com/), resulting fastq outputs were adapter trimmed and low-quality reads discarded using cutadapt (version 2.5)[14]. Cleaned fastq files were mapped against GCRh37p5 using BWA[2] (version 0.7.12), and sample targets were extracted from the resulting BAM file using SAMtools (version 0.1.19)[15]. Aligned ONT reads were phased using an in-house tool, with frequencies and pre/post zygosity calls affirmed via IGV and principal component analysis using the available exome sequence data for probands and parents to support the ONT data.

CNV analysis

CNV calling was performed on our trio-based exome data with a custom GATK4-based pipeline (Mikulasova et al., in preparation). This workflow exploits the GATK4 sequence read-depth normalization[16] and a custom R based segmentation and visualization[17]. Parental samples from the trios under examination were used as controls for the normalization step. The CNVs detected were annotated using AnnotSV (https://lbgi.fr/AnnotSV/)[18]. CNVs present in more than 1% of the samples of the Database of Genomic Variations or present in more than 10% of the patients were excluded from the analysis. The remaining rare deletions and duplications were individually inspected through the genomic profiles and detailed Log2Ratio plots generated by the workflow. Only CNVs involving more than 2 exons were further considered to minimize the inclusion of false positives, and we selected 2 CNVs present in the probands but absent in their parents for further validation. Validation was performed with the whole genome Illumina Infinium CytoSNP-850K v1.1 microarray platform for the larger deletion (chromosome 11) and a gene specific TaqMan Copy Number assay (designed for *NXT2*) was exploited to validate the smaller one.

Functional enrichment

To evaluate the intolerance of each gene for loss-of-function (LoF) mutations, we used the probability of LoF intolerance (pLI) score, based on data from the Genome Aggregation Database (GnomAD)[19] containing genetic data from 141,456 individuals. We computed the likelihood of the observed median pLI score of each gene (LoF in controls) set compared to the expected median pLI based on the method described in Lelieveld et al[20]. Instead of using the complete set of 18,226 pLI annotated genes to obtain expected median pLi scores, we used a set of 2,766 coding DNMs in 1,941 control individuals to correct for the gene-specific mutation rate downloaded from the denovo-db version 1.6.1 (http://denovo-db.gs.washington.edu/denovo-db/)[21]. To evaluate the impact of the *de novo* missense mutations to each gene, we used missense Z scores calculated by GnomAD[19,22] to predict the tolerance of each gene to variation. The presence of missense mutations in intolerant genes was compared between predicted pathogenic and benign using the Chi-Squared test, adjusted for multiple testing using Bonferroni correction. To predict the affected protein function and the potential role in disease, we evaluated the interactions between the genes with a DNM using STRING version 11[23].

Additional Cohorts of Infertile Men

The strongest candidate genes with DNMs were further investigated in exome data from four additional cohorts of infertile men. For the Italian cohort of 48 patients with NOA, exome sequencing was carried out as a service by Macrogen Inc. (Republic of Korea) utilizing the Agilent SureSelect_V6 enrichment and a NovaSeq6000. The German Male Reproductive Genomics (MERGE) study comprised exome data of 887 men with azoo-, crypto-, or severe oligozoospermia. Known causes for male infertility like chromosomal aberrations and microdeletions of the AZF region were excluded in advance. Whole exome sequencing was performed as previously described[24]. The 88 patients diagnosed with male infertility participating in the Geisinger-Regeneron DiscovEHR collaboration were selected from deidentified EHR information using the ICD-10CM code N46 which refers to 'Male Infertility' including oligospermia, azoospermia, other male infertility and male infertility unspecified. All patients were sequenced at the Regeneron Genetics Center (RGC) as previously described[25]. In brief, 1ug of genomic DNA per sample was used for targeted exome capture using the NimbleGen VCRome 2.1 or the IDT XGen reagents. Captured libraries were sequenced on the Illumina HiSeq 2500 platform with v4 chemistry using paired-end 75 bp reads. Exome sequencing was performed such that >85% of the bases were covered at 20x or greater. Raw sequence reads were mapped and aligned to the GRCh38/hg38 human genome reference assembly using BWA-mem[2] and single nucleotide and indel variants and genotypes were called using GATK's HaplotypeCaller[5]. The Genetics of Male Infertility INitiative (GEMINI) is a multicenter study funded by the United States NIH. The GEMINI project performed whole-exome sequencing on 1,011 unrelated men diagnosed with spermatogenic failure, the vast majority with unexplained NOA (Nagrinaja, et al. in preparation). Sequencing of genomic DNA was performed at the McDonnell Genome Institute of Washington University in St. Louis, MO, USA, using an in-house exome targeting reagent capturing 39.1 Mb of exome and 2 x 150 bp paired-end sequencing on Illumina HiSeq 4000. Following sample QC, a final cohort of 924 men were analyzed as part of the current study. DiscovEHR participants were sequenced at the Regeneron Genetics Center (RGC) as previously described[25]. In brief, 1ug of genomic DNA per sample was used for targeted exome capture using the NimbleGen VCRome 2.1 or the IDT XGen reagents. Captured libraries were sequenced on the Illumina HiSeq 2500 platform with v4 chemistry using paired-end 75 bp reads. Exome sequencing was performed such that >85% of the bases were covered at 20x or greater. Raw sequence reads were mapped and aligned to the GRCh38/hg38 human genome reference assembly using BWA-mem[2] and single nucleotide and indel variants and genotypes were called using GATK's HaplotypeCaller[5].

Genetic variants identified within the 33 candidate genes were extracted from each exome dataset. Consistently with our filtering method described above variants with <10 reads and/or less than 15% reads containing the mutation were discarded. To minimize discrepancies between genomic positions and annotation, genomic coordinated were recalculated to the GCRh37/hg19 and fully reannotated with VEP[6]. Following annotations, variants with allele frequency >1% in gnomAD were discarded to focus only on rare variants. Pathogenicity predictions based on SIFT[7], MutationTaster[8] and PolyPhen[9] were then used to exclude benign variants, all remaining variants were classified according to ACMG guidelines[10]. Like before, variants found in one or more fathers in our control cohort of verified fertile parents were removed from further consideration.

Burden testing

Having identified several likely pathogenic rare loss-of-function and missense mutations in these 35 genes we performed a gene-based burden test to compare the combined data in all cohorts of infertile men with the control cohort of fertile fathers. The proportion of individuals with pathogenic variants was statistically evaluated using Fisher's Exact test, adjusted for multiple testing following Bonferroni correction. Similarly, a gene-based burden test was performed to compare fertile fathers with fertile mothers from the control cohort of verified fertile parents to investigate whether any of the sexes predominantly carried a greater number of rare pathogenic mutations.

Data Availability

Raw and processed data is available under controlled access and requires a Data Transfer Agreement from the European Genome-Phenome Archive (EGA) repository: EGAS00001004945.
