## Supplementary material for "A *de novo* paradigm for male infertility": Suplementary Figures 1-8

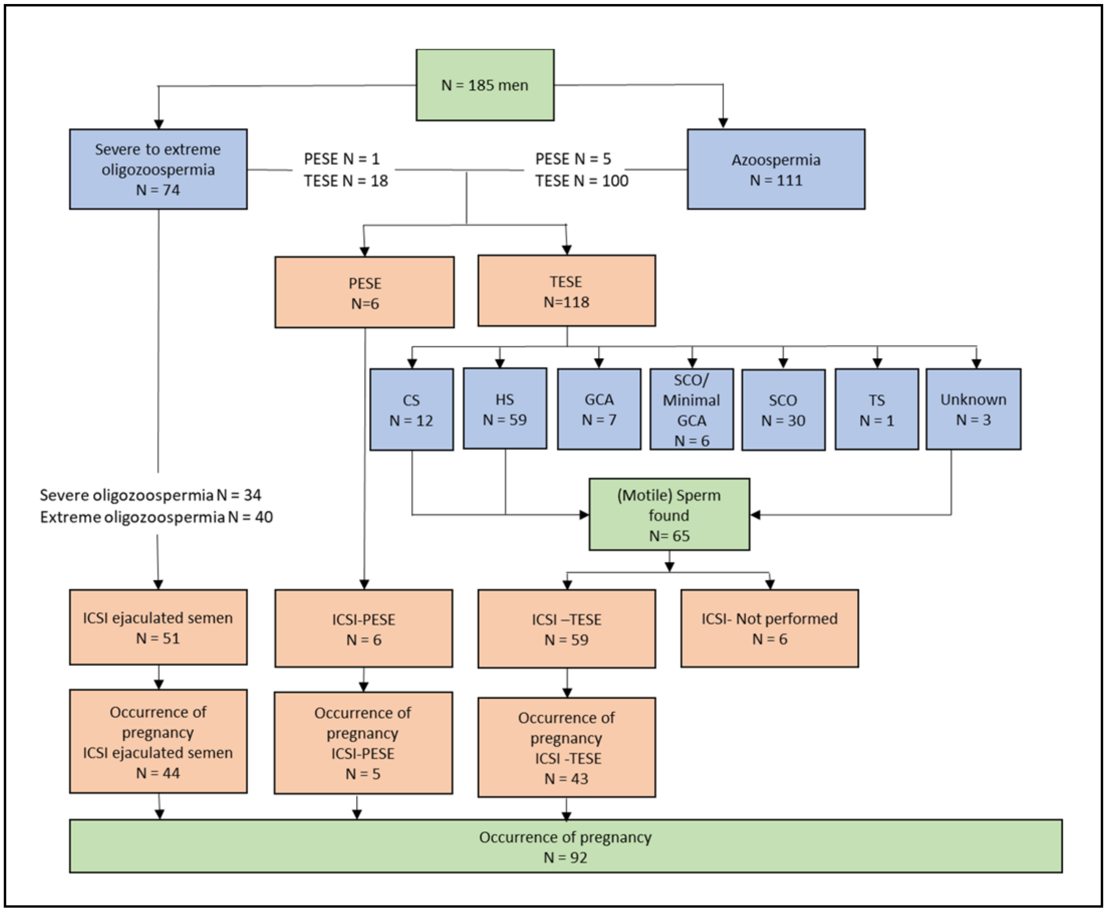


**Supplementary Figure 1**: **Flow chart of diagnostic and clinical outcome**. CS = complete spermatogenesis, HS =hypospermatogenesis, GCA = germ cell arrest, SCO = Sertoli cell only, ICSI= intracytoplasmic sperm injection, PESA=percutaneous sperm aspiration, TESE= testicular sperm extraction. TS= tubular shadows

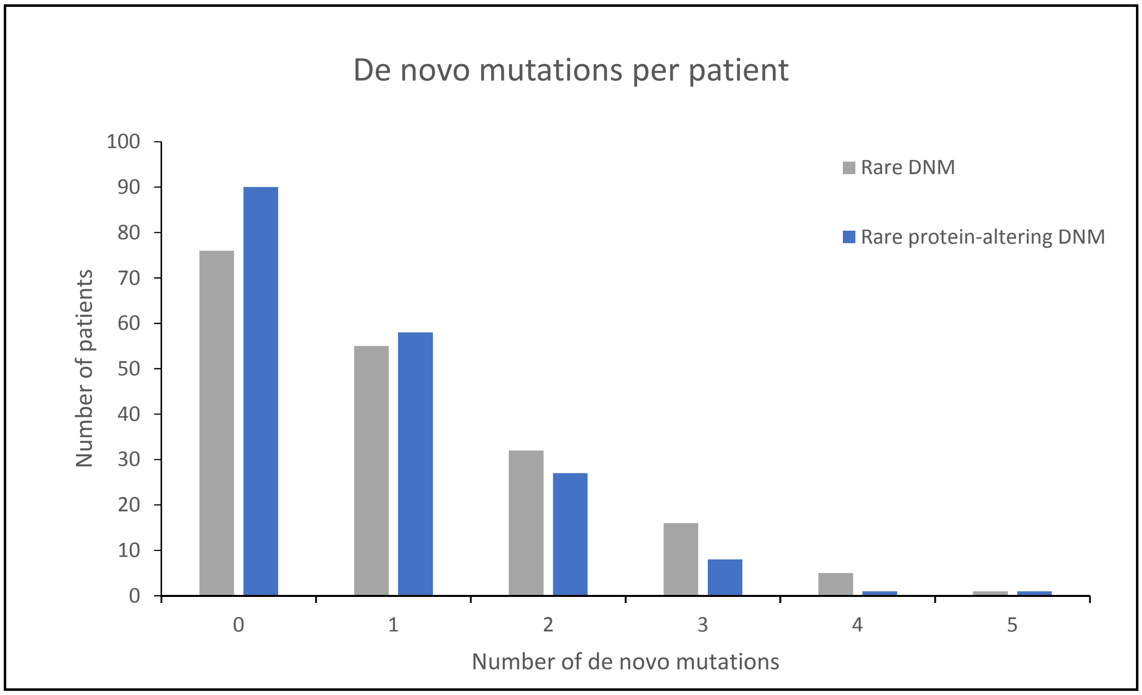


**Supplementary Figure 2: Distribution of 192 *de novo* mutations in 185 patients.**

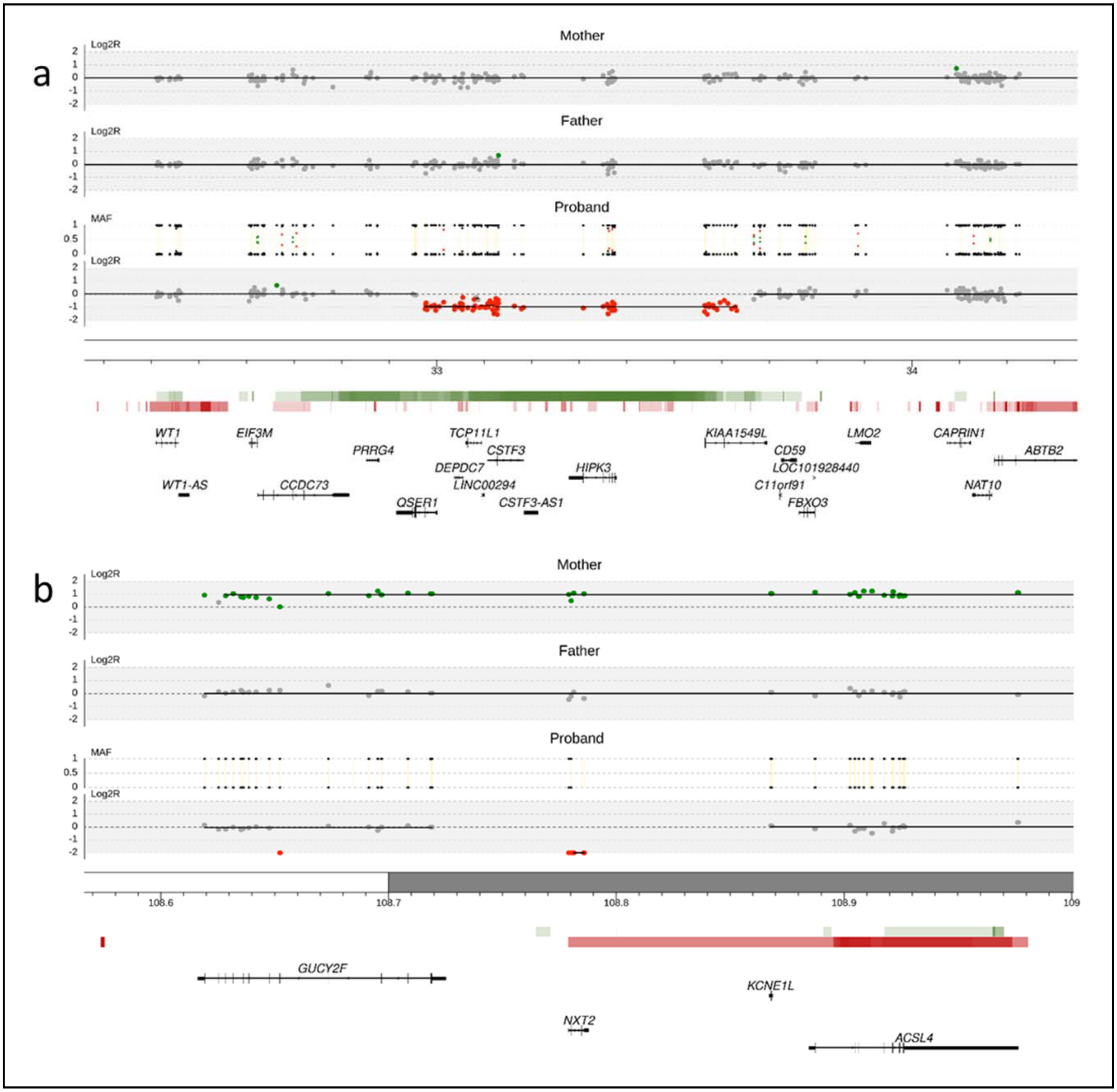


**Supplementary Figure 3: De novo Copy Number Variations (CNVs) identified in infertile men**. **a,** De novo deletion of ~656kb (chr11:32975325-33631588) identified in patient Proband_066 affecting genes *QSER1* (partially*), DEPDC7, TCP11L1, LINC00294, CSTF3, HIPK3* and *KIAA1549L* (partially). **b,** Deletion of ~6kb (chrX:108779109-108785919) identified in Proband_039 affecting gene *NXT2*. Log2Ratio tracks show the number of alleles in the specific region of chromosome in mother, father and proband, which also include the Minor Allele Frequency (MAF) plot showing SNP zygosity. Below the Log2R plots are the cytobands of the specific region of the chromosome. Horizontal red and green bars indicate respectively the deletions and duplications in the Database of Genomic Variants (DGV). The more intense the colour, the more CNVs are present in the region. Below the DGV track, RefSeq genes are indicated.

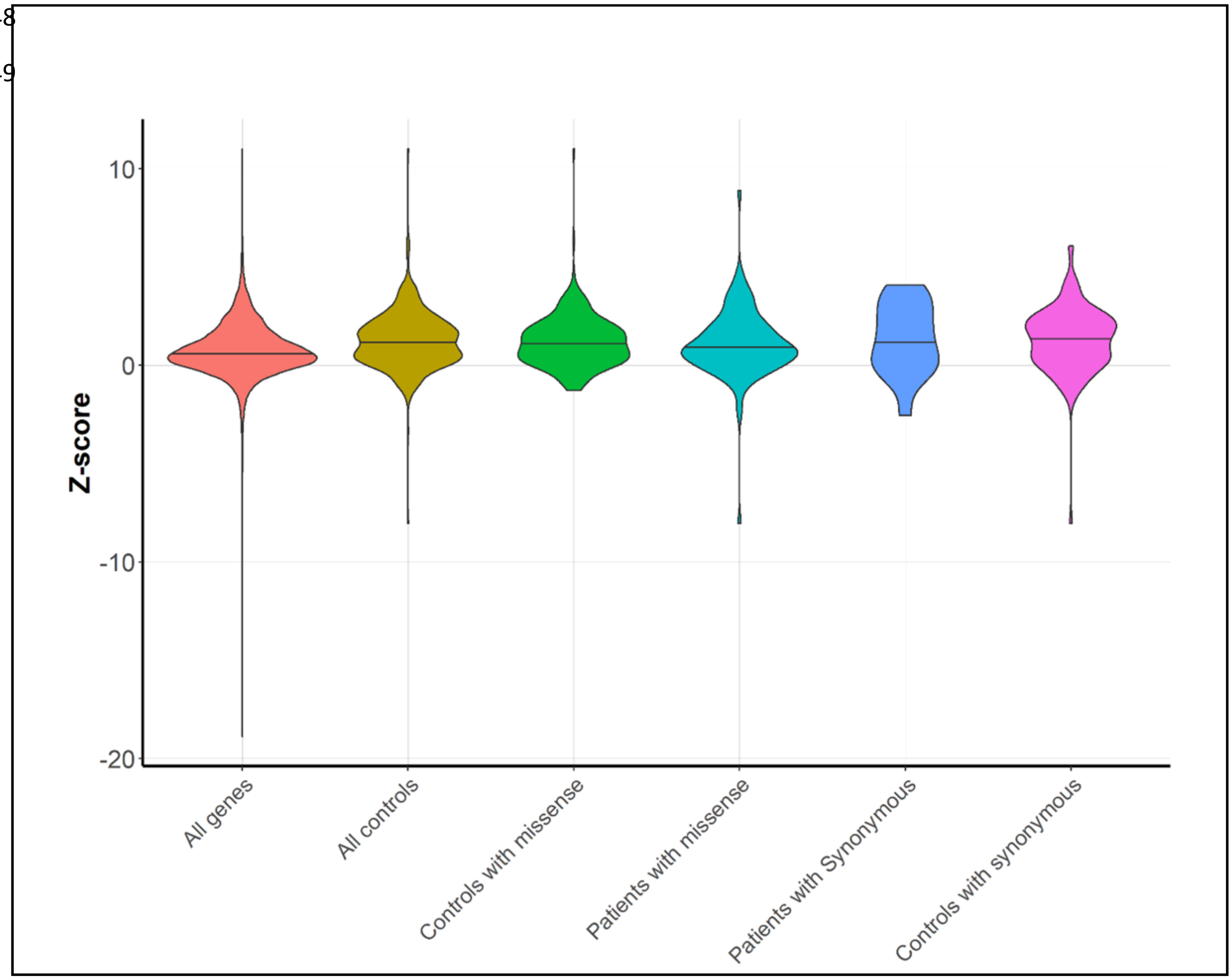


**Supplementary Figure 4: Analysis of the intolerance to missense variants for DNM genes.** Violin plots represent the distribution of the z-scores of all genes in gnomAD, all genes affected by missense DNMs and all synonymous DNM in this study and in a control population (http://denovo-db.gs.washington.edu/denovo-db/). The higher the z-score, the more intolerant the gene is to missense variants. Comparison between overall missense DNMs in our study and control populations shows no significant difference.

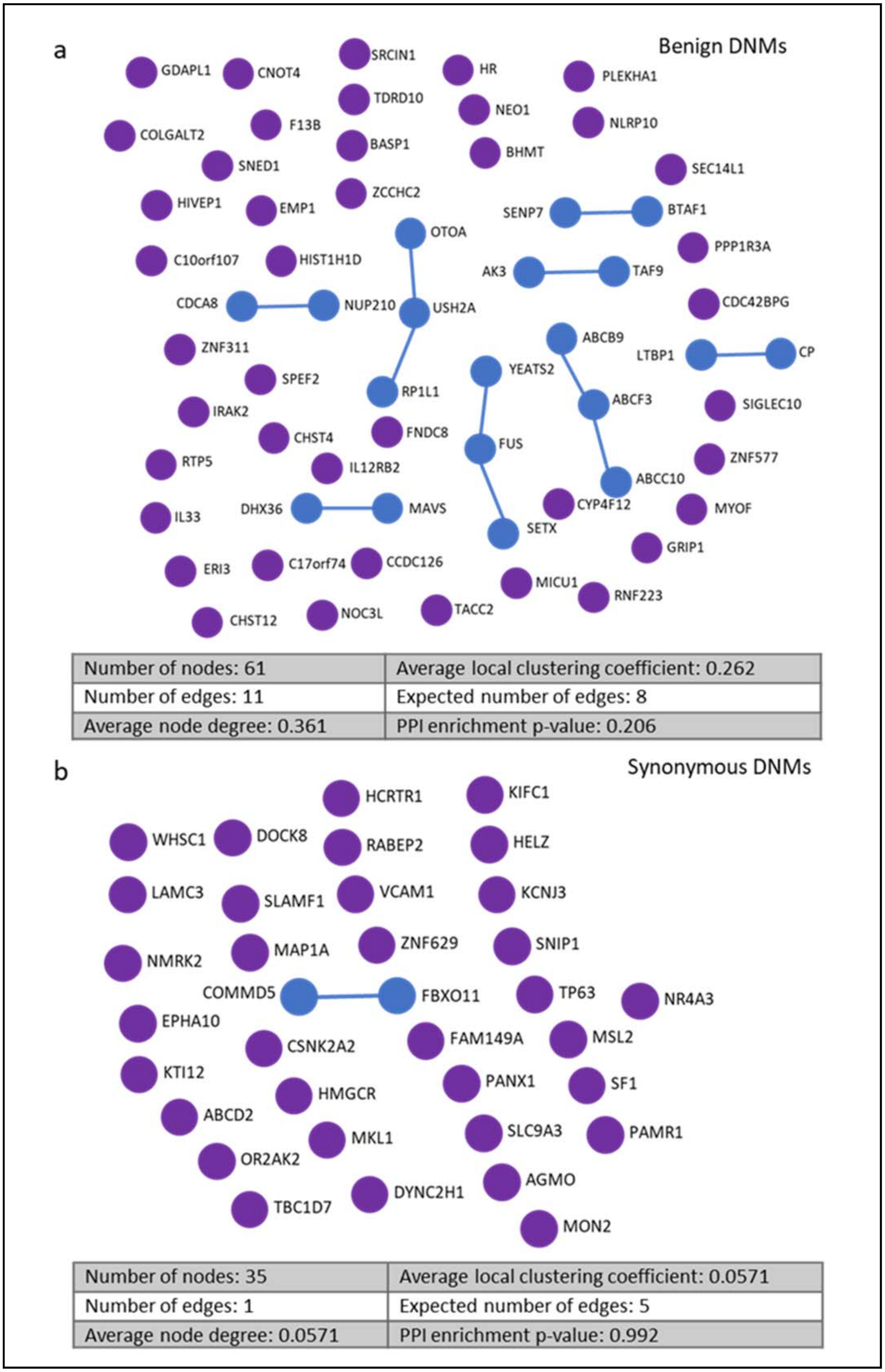


**Supplementary Figure 5: Protein-protein interactions between synonymous DNM.** a) A protein-protein interaction analysis was performed on all protein-coding benign DNM (n=61) which classified as benign in 2/3 of the pathogenicity scores. No significant interaction is seen between the different proteins (PPI enrichment p-value= 0.206). b) A protein-protein interaction analysis was performed on all synonymous DNM which are known to not affect the gene. Here no significant interaction is seen with fewer edges than would be expected between this number of genes (PPI enrichment p-value 0.992).

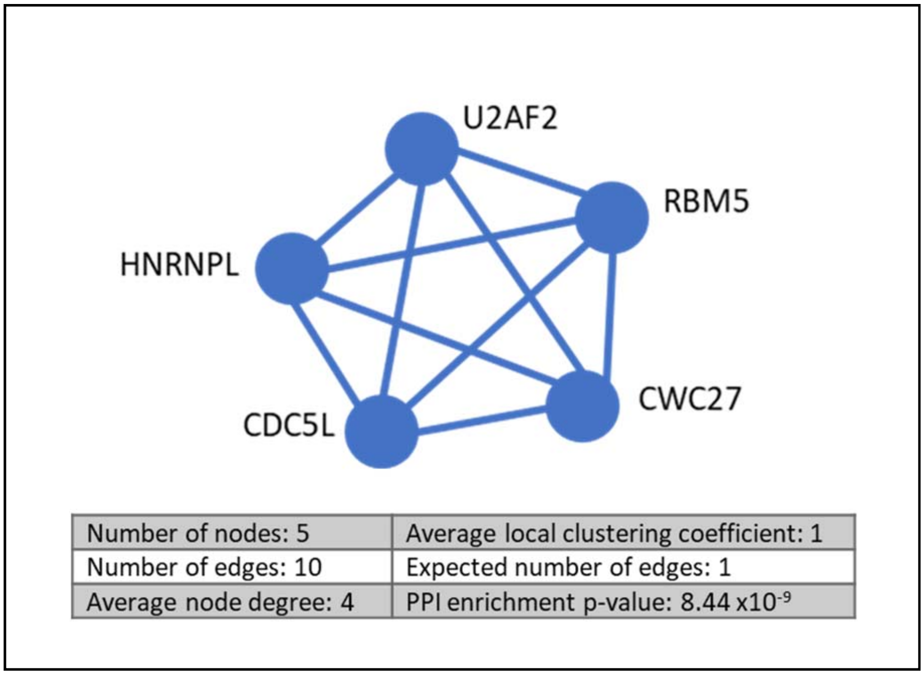


**Supplementary Figure 6: Central module of protein-protein interaction network.** The 5 proteins: *RBM5*, *HNRNPL*, *CWC27*, *CDC5L* and *U2AF2* all interact highly with one another (PPI enrichment p-value = $8.44 \times 10^{-9}$) and are all seen to be involved in the biological process of mRNA splicing, via the spliceosome (False discovery rate: $1.72 \times 10^{-7}$).

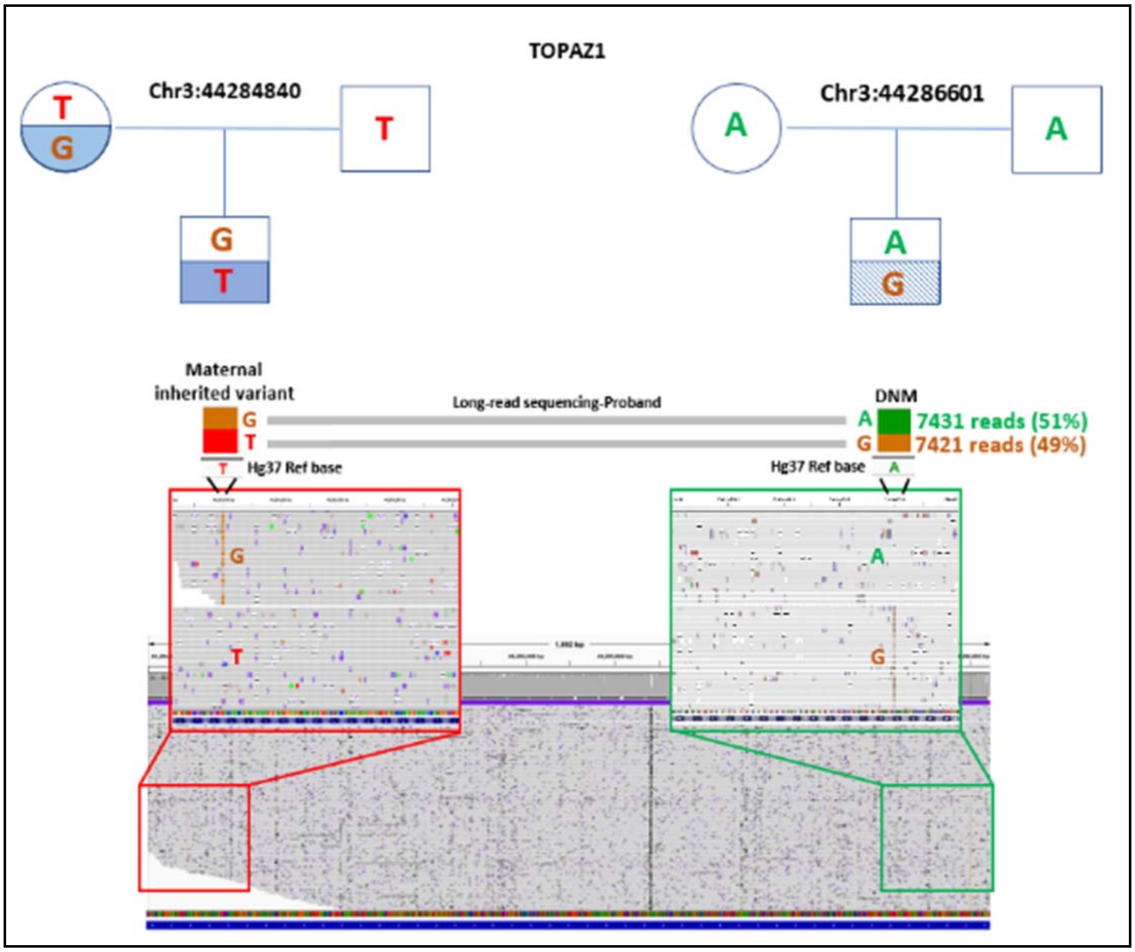


**Supplementary Figure 7: Pedigree and IGV illustration of *TOPAZ1*.** Position and bases of the maternally inherited variant and the DNM that were identified in *TOPAZ1* are depicted by the pedigree chart. Using long-read targeted sequencing, the ~2000 bp region that these variants span is sequenced and visualized in IGV. 14852 reads are phased, showing a 51/49 percentage split. The DNM is identified as paternal in origin and thus on the alternate allele to the inherited variant.

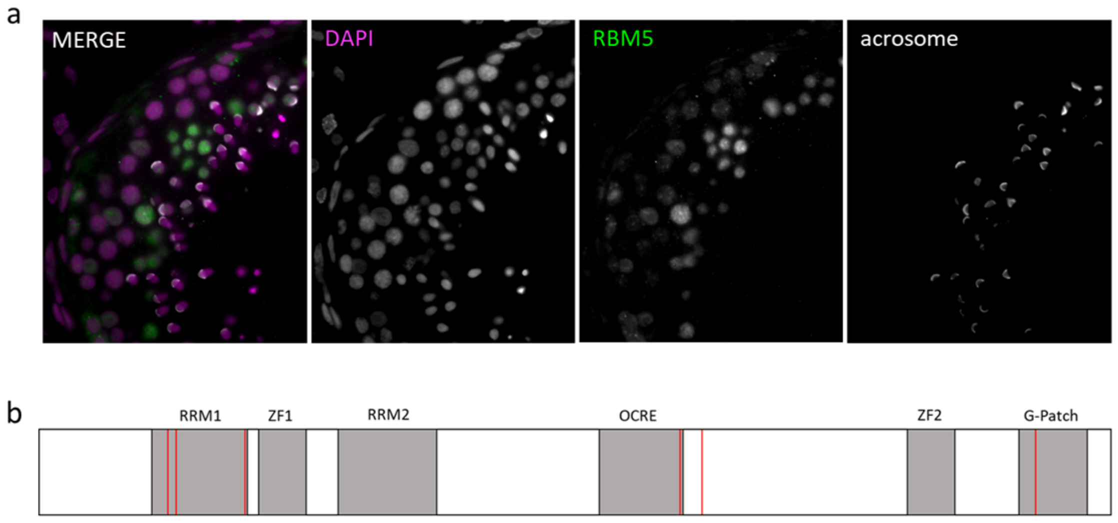


c

| Cohort | Sample_ID | Genetic Coordinates (GRCh37) | Refseq ID | HGVS |
|---|---|---|---|---|
| Regeneron DiscovEHR Cohort of Infertile Men | 742146 | chr3:50137439 | NM_005778.4:c.364G>A | p.Glu122Lys |
| MERGE Cohort of Infertile Men | M248 | chr3:50137446 | NM_005778.4:c.371C>T | p.Pro124Leu |
| NIJ/NCL Cohort of Patient-Parent Trios (DNM) | Proband_108 | chr3:50140556 | NM_005778.4:c.524A>G | p.Tyr175Cys |
| NIJ/NCL Cohort of Infertile Men (Singleton) | Proband00524 | chr3:50150877 | NM_005778.4:c.1517T>A | p.Val506Glu |
| MERGE Cohort of Infertile Men | M2013 | chr3:50150955 | NM_005778.4:c.1595C>T | p.Pro532Leu |
| NIJ/NCL Cohort of Infertile Men (Singleton) | Proband00282 | chr3:50154750 | NM_005778.4:c.2260A>G | p.Met754Val |

**Supplementary Figure 8: *RBM5* pathogenic mutations found in multiple infertile men from 4 different international cohorts** a) Localisation of *RBM5* in human testis. DAPI in magenta, *RBM5* in green and the acrosome in white. *RBM5* is expressed in most stages of germ cell development albeit at different levels. Expression in Sertoli cells is also observed. b) Schematic representation of *RBM5* protein domains and the location of rare pathogenic mutations found in infertile males. c) Details of pathogenic variants found in 6 infertile males.
