## Supplementary Table 1 for "A *de novo* paradigm for male infertility"

| Patient ID | Origin | Semen analysis | *Gene* | Chromosome coordinates (GRCh37) | Refseq ID | HGVS | Pathogenicity prediction* | Expressed in testis |
|---|---|---|---|---|---|---|---|---|
| Proband_005 | Netherlands | Azoospermia | *CDK5RAP2* | chr9:123215805 | NM_018249:c.2722C>T | p.Arg908Trp | SP | Yes, not enhanced |
| Proband_006 | Netherlands | Azoospermia | *ATP1A1* | chr1:116930014 | NM_000701:c.291del | p.Phe97LeufsTer44 | N/A | Yes, not enhanced |
| | | | *TLN2* | chr15:63029134 | NM_015059:c.3416G>A | p.Gly1139Glu | MP | Yes, not enhanced |
| | | | *HUWE1* | chrX:53589090 | NM_031407:c.7314_7319del | p.Glu2439_Glu2440del | N/A | Yes, not enhanced |
| Proband_008 | Netherlands | Azoospermia | *ABCC10* | chr6:43417749 | NM_001198934:c.4399C>T | p.Arg1467Cys | - | Yes, not enhanced |
| | | | *CP* | chr3:148927135 | NM_000096c.644G>A | p.Arg215Gln | - | Not expressed |
| Proband_010 | Netherlands | Azoospermia | *FUS* | chr16:31196402 | NM_004960:c.678_686del | p.Gly229_Gly231del | N/A | Yes, not enhanced |
| | | | *LTBP1* | chr2:33246090 | NM_206943:c.680C>G | p.Ser227Trp | P | Yes, not enhanced |
| Proband_012 | Netherlands | Extreme oligozoospermia | *RP1L1* | chr8:10480174 | NM_178857:c.538G>A | p.Ala180Thr | SP | Yes, not enhanced |
| Proband_013 | Netherlands | Azoospermia | *ERG* | chr21:39755563 | NM_182918:c.1202C>T | p.Pro401Leu | MP | Yes, not enhanced |
| Proband_017 | Netherlands | Azoospermia | *CDC5L* | chr6:44413480 | NM_001253:c.2180G>A | p.Arg727His | SMP | Yes, not enhanced |
| Proband_019 | Netherlands | Azoospermia | *ABLIM1* | chr10:116205100 | NM_002313:c.1798C>T | p.Arg600Trp | SMP | Yes, not enhanced |
| Proband_020 | Netherlands | Azoospermia | *CCDC126* | chr7:23682709 | NM_001253:c.2180G>A | p.Thr133Met | - | Yes, enhanced expression in testis |
| | | | *RASEF* | chr9:85607885 | NM_152573:c.1976G>A | p.Arg659His | SMP | Yes, not enhanced |
| Proband_022 | Netherlands | Azoospermia | *APC2* | chr19:1453118 | NM_005883:c.118G>A | p.Glu40Lys | MP | Yes, not enhanced |
| Proband_025 | Netherlands | Azoospermia | *NEO1* | chr15:73575428 | NM_002499:c.3386G>A | p.Arg1129His | SMP | Yes, not enhanced |
| Proband_028 | Netherlands | Extreme oligozoospermia | *OR5P3* | chr11:7846930 | NM_153445:c.590T>A | p.Ile197Lys | SP | No expression |
| Proband_030 | Netherlands | Extreme oligozoospermia | *SIKE1* | chr1:115323119 | NM_025073:c.110A>C | p.His37Pro | MP | Yes, not enhanced |
| | | | *TRAF7* | chr16:2218149 | NM_032271:c.211C>T | p.Arg71Trp | SMP | Yes, not enhanced |
| Proband_033 | Netherlands | Azoospermia | *KRT33B* | chr17:39521752 | NM_002279:c.642C>G | p.Asp214Glu | SP | No expression |
| | | | *SENP7* | chr3:101212750 | NM_020654:c.153C>G | p.Phe51Leu | - | Yes, not enhanced |
| | | | *ATP8A1* | chr4:42626573 | NM_006095:c.343A>G | p.Lys115Glu | SMP | Yes, not enhanced |

| | | | | | | | | |
|---|---|---|---|---|---|---|---|---|
| Proband_038 | Netherlands | Azoospermia | *NOC3L* | chr10:96100055 | NM_022451:c.1758A>C | p.Lys586Asn | M | Yes, not enhanced |
| Proband_039 | Netherlands | Azoospermia | *NXT2* | chrX:108779109:108785919 | N/A | N/A | N/A | Yes, not enhanced |
| Proband_041 | Netherlands | Azoospermia | *ASIC5* | chr4:156763436 | NM_017419:c.932G>A | p.Ser311Asn | SM | Not expressed |
| Proband_042 | Netherlands | Azoospermia | *PLCL1* | chr2:198966024 | NM_006226:c.2935C>T | p.Arg979Trp | SMP | Yes, not enhanced |
| | | | *DNAJC2* | chr7:102957321 | NM_014377:c.1383A>T | p.Leu461Phe | SMP | Yes, enhanced expression in testis |
| | | | *AK3* | chr9:4722547 | NM_016282:c.230A>G | p.His77Arg | - | Yes, not enhanced |
| Proband_043 | Netherlands | Azoospermia | *IL33* | chr9:6251221 | NM_033439:c.299G>A | p.Gly100Glu | - | Yes, not enhanced |
| Proband_044 | Netherlands | Azoospermia | *PRDM16* | chr1:3328833 | NM_022114:c.2072A>T | p.Asp691Val | SMP | Yes, not enhanced |
| | | | *PPP1R7* | chr2:242099831 | NM_002712:c.523_527del | p.Lys175GlnfsTer2 | N/A | Yes, not enhanced |
| Proband_045 | Netherlands | Azoospermia | *EVC* | chr4:5743515 | NM_153717:c.775C>T | p.Gln259Ter | N/A | Yes, not enhanced |
| | | | *BHMT* | chr5:78426892 | NM_001713:c.1174C>G | p.Gln392Glu | M | Yes, not enhanced |
| Proband_048 | Netherlands | Extreme oligozoospermia | *MCM6* | chr2:136624195 | NM_005915:c.719A>G | p.Asp240Gly | SMP | Yes, not enhanced |
| Proband_049 | Netherlands | Azoospermia | *ATP8B4* | chr15:50168651 | NM_024837:c.2851A>G | p.Asn951Asp | SMP | Yes, not enhanced |
| | | | *ZNF577* | chr19:52376320 | NM_032679:c.923A>G | p.Tyr308Cys | P | Yes, not enhanced |
| | | | *NUP210* | chr3:13359251 | NM_024923:c.5594A>G | p.Asn1865Ser | - | Yes, not enhanced |
| Proband_050 | Netherlands | Extreme oligozoospermia | *HIST1H1D* | chr6:26234816 | NM_005320:c.346G>C | p.Glu116Gln | M | Yes, not enhanced |
| Proband_051 | Netherlands | Extreme oligozoospermia | *FNDC8* | chr17:33448840 | NM_017559:c.128G>A | p.Arg43Gln | S | Yes, enhanced expression in testis |
| Proband_052 | Netherlands | Severe oligozoospermia | *SOGA1* | chr20:35438426 | NM_080627:c.2542C>T | p.Arg848Ter | N/A | Yes, not enhanced |
| Proband_053 | Netherlands | Extreme oligozoospermia | *CD81* | chr11:2417877 | NM_004356:c.581T>C | p.Ile194Thr | SMP | Yes, not enhanced |
| Proband_055 | Netherlands | Azoospermia | *OSBPL3* | chr7:24874131 | NM_015550:c.1720C>T | p.Gln574Ter | N/A | Yes, not enhanced |
| Proband_057 | Netherlands | Extreme oligozoospermia | *ABCB9* | chr12:123430670 | NM_019625:c.1153G>A | p.Glu385Lys | M | Yes, enhanced expression in testis |
| Proband_060 | Netherlands | Azoospermia | *IL12RB2* | chr1:67861761 | NM_001559:c.2578C>G | p.Leu860Val | P | Yes, not enhanced |
| | | | *TOPAZ1* | chr3:44286601 | NM_001145030:c.2603A>G | p.Gln868Arg | SP | Yes, enhanced expression in testis |
| Proband_061 | Netherlands | Azoospermia | *SEC14L1* | chr17:75208114 | NM_001143999:c.1694A>G | p.Tyr565Cys | S | Yes, not enhanced |
| Proband_062 | Netherlands | Extreme oligozoospermia | *FOXF2* | chr6:1391182 | NM_001452:c.1000A>G | p.Thr334Ala | SM | Yes, not enhanced |
| Proband_063 | Netherlands | Extreme oligozoospermia | *CNOT4* | chr7:135048789 | NM_001190849:c.1657A>G | p.Met553Val | M | Yes, not enhanced |
| Proband_064 | Netherlands | Severe oligozoospermia | *C9orf50* | chr9:132374702 | NM_199350:c.1220C>T | p.Ser407Leu | SP | Yes, enhanced expression in testis |

| | | | | | | | | |
|---|---|---|---|---|---|---|---|---|
| Proband_066 | Sint Maarten (Caribbean) | Extreme oligozoospermia | *HIPK3* | chr11:32975325:33631588 | N/A | N/A | N/A | Yes, not enhanced |
| | | | *QSER1* | | N/A | N/A | | Yes, not enhanced |
| | | | *DEPDC7* | | N/A | N/A | | Yes, enhanced expression in testis |
| | | | *TCP11L1* | | N/A | N/A | | Yes, not enhanced |
| | | | *CSTF3* | | N/A | N/A | | Yes, not enhanced |
| | | | *KIAA1549L* | | N/A | N/A | | Yes, not enhanced |
| | | | *TDRD10* | chr1:154493890 | NM_182499:c.304G>A | p.Val102Met | P | Yes, enhanced expression in testis |
| | | | *CWC27* | chr5:64077814 | NM_005869:c.206C>G | p.Thr69Ser | MP | Yes, not enhanced |
| Proband_073 | Netherlands | Extreme oligozoospermia | *INO80* | chr15:41372056 | NM_017553:c.974C>T | p.Ala325Val | SM | Yes, not enhanced |
| Proband_074 | Netherlands | Severe oligoasthenozoospermia | *ZNF709* | chr19:12575362 | NM_152601:c.1375dup | p.Ser458ArgfsTer10 | N/A | Yes, enhanced expression in testis |
| | | | *EMILIN1* | chr2:27305208 | NM_007046:c.769G>A | p.Glu257Lys | MP | Yes, not enhanced |
| | | | *WDR17* | chr4:177067235 | NM_170710:c.1619G>A | p.Gly540Glu | MP | Yes, not enhanced |
| | | | *ZNF311* | chr6:28963503 | NM_001010877:c.1276C>T | p.Arg426Trp | S | Yes, not enhanced |
| Proband_076 | Netherlands | Azoospermia | *STARD10* | chr11:72466763 | NM_006645:c.610_612del | p.Ser204del | N/A | Yes, not enhanced |
| | | | *GREB1L* | chr18:19019514 | NM_001142966:c.868_872del | p.Gly290CysfsTer19 | N/A | Yes, enhanced expression in testis |
| Proband_077 | Netherlands | Extreme oligozoospermia | *MSH5* | chr6:31721100 | NM_172165:c.887_888del | p.His296ArgfsTer90 | N/A | Yes, not enhanced |
| Proband_079 | Netherlands | Azoospermia | *ILVBL* | chr19:15226717 | NM_006844:c.1558G>A | p.Gly520Arg | SMP | Yes, not enhanced |
| Proband_080 | Netherlands | Extreme oligozoospermia | *HOXA1* | chr7:27134363 | NM_005522:c.704A>C | p.Asn235Thr | SMP | Yes, not enhanced |
| Proband_081 | Netherlands | Azoospermia | *ZFHX4* | chr8:77763486 | NM_024721:c.4331del | p.Leu1444TrpfsTer8 | N/A | Yes, enhanced expression in testis |
| Proband_083 | Netherlands | Azoospermia | *F13B* | chr1:197026291 | NM_001994:c.1023C>A | p.Phe341Leu | - | Not expressed |
| | | | *HNRNPL* | chr19:39329152 | NM_001533:c.1442G>A | p.Arg481Gln | SMP | Yes, not enhanced |
| Proband_085 | Netherlands | Severe oligoasthenozoospermia | *NLRP10* | chr11:7981967 | NM_176821:c.1192G>A | p.Asp398Asn | - | Not expressed |
| Proband_087 | Netherlands | Severe oligoasthenozoospermia | *LEO1* | chr15:52252183 | NM_138792:c.1073T>G | p.Ile358Arg | SMP | Yes, not enhanced |
| Proband_088 | Netherlands | Extreme oligozoospermia | *GDAP1L1* | chr20:42893167 | NM_024034:c.728C>T | p.Ala243Val | P | Yes, not enhanced |
| Proband_095 | Netherlands | Azoospermia | *MPRIP* | chr17:17062191 | NM_015134:c.1921C>T | p.His641Tyr | MP | Yes, not enhanced |
| | | | *SORCS2* | chr4:7728558 | NM_020777:c.2797G>A | p.Asp933Asn | SM | Yes, not enhanced |
| Proband_097 | Netherlands | Azoospermia | *TENM2* | chr5:167642269 | NM_001122679.2:c.4046delC | p.Pro1349ArgfsTer6 | N/A | Yes, not enhanced |
| Proband_101 | Netherlands | Extreme oligozoospermia | *CHST12* | chr7:2472611 | NM_001243794:c.337C>T | p.Arg113Cys | - | Not expressed |

| | | | | | | | | |
|---|---|---|---|---|---|---|---|---|
| Proband_102 | Netherlands | Extreme oligozoospermia | *HR* | chr8:21973239 | NM_005144:c.3544G>A | p.Val1182Met | SMP | Yes, not enhanced |
| | | | *SMC2* | chr9:106885442 | NM_006444:c.2186T>C | p.Leu729Ser | MP | Yes, not enhanced |
| Proband_106 | Netherlands | Azoospermia | *TACC2* | chr10:123976284 | NM_206862:c.7487C>T | p.Pro2496Leu | SMP | Yes, not enhanced |
| Proband_108 | Netherlands | Severe oligoasthenozoospermia | *CYP4F12* | chr19:15794526 | NM_023944:c.871G>A | p.Ala291Thr | - | Not expressed |
| | | | *RBM5* | chr3:50140556 | NM_005778:c.524A>G | p.Tyr175Cys | SMP | Yes, not enhanced |
| | | | *TAF9* | chr5:68660785 | NM_003187:c.777_779del | p.Asp260del | - | Yes, not enhanced |
| Proband_115 | Netherlands | Severe oligoasthenozoospermia | *CDC42BPG* | chr11:64603286 | NM_017525:c.1706C>T | p.Thr569Met | - | Yes, not enhanced |
| | | | *RPA1* | chr17:1756424 | NM_002945:c.302T>C | p.Val101Ala | SMP | Yes, not enhanced |
| Proband_116 | Netherlands | Severe oligoasthenozoospermia | *FBXO5* | chr6:153293449 | NM_012177:c.1046_1049dup | p.Asp350GlufsTer9 | N/A | Yes, not enhanced |
| Proband_117 | Netherlands | Azoospermia | *FLNC* | chr7:128477754 | NM_001458:c.914C>T | p.Ala305Val | SMP | Yes, not enhanced |
| Proband_118 | Netherlands | Azoospermia | *CDCA8* | chr1:38166170 | NM_001256875:c.400C>T | p.Arg134Cys | S | Yes, enhanced expression in testis |
| Proband_119 | Netherlands | Severe oligoasthenozoospermia | *ZCCHC2* | chr18:60242391 | NM_017742:c.3077A>G | p.Asn1026Ser | - | Yes, not enhanced |
| Proband_121 | Netherlands | Azoospermia | *AMPD2* | chr1:110168336 | NM_139156:c.194A>T | p.Glu65Val | SMP | Yes, not enhanced |
| | | | *HIVEP1* | chr6:12163643 | NM_002114:c.7106C>T | p.Pro2369Leu | - | Yes, not enhanced |
| | | | *SPEF2* | chr5:35792492 | NM_024867:c.4498C>G | p.Leu1500Val | - | Yes, not enhanced |
| Proband_122 | Netherlands | Azoospermia | *SIGLEC10* | chr19:51919175 | NM_033130:c.1001G>A | p.Arg334Gln | P | Not expressed |
| Proband_124 | Netherlands | Severe oligoasthenozoospermia | *PPP1R3A* | chr7:113518521 | NM_002711:c.2626T>G | p.Phe876Val | P | Not expressed |

| | | | | | | | | |
|---|---|---|---|---|---|---|---|---|
| Proband_125 | Netherlands | Azoospermia | *CHST4* | chr16:71571634 | NM_001166395:c.1054G>A | p.Asp352Asn | - | Not expressed |
| | | | *STXBP2* | chr19:7711198 | NM_006949:c.1420C>T | p.Arg474Cys | SMP | Yes, not enhanced |
| Proband_126 | Netherlands | Severe oligoasthenozoospermia | *ABCF3* | chr3:183907504 | NM_018358:c.1273C>T | p.Arg425Cys | SMP | Yes, not enhanced |
| Proband_127 | Netherlands | Extreme oligozoospermia | *TMEM62* | chr15:43441280 | NM_024956:c.797C>G | p.Pro266Arg | SMP | Yes, not enhanced |
| | | | *U2AF2* | chr19:56170622 | NM_007279:c.100_105dup | p.Ser34_Arg35dup | N/A | Yes, not enhanced |
| Proband_128 | Netherlands | Azoospermia | *HELZ2* | chr20:62195532 | NM_001037335:c.4643C>T | p.Thr1548Met | SP | Yes, not enhanced |
| Proband_129 | Netherlands | Azoospermia | *FIZ1* | chr19:56104069 | NM_032836:c.1235_1237del | p.Lys412del | N/A | Yes, not enhanced |
| Proband_130 | Netherlands | Azoospermia | *MAVS* | chr20:3846631 | NM_020746:c.1460C>T | p.Ala487Val | - | Yes, not enhanced |
| | | | *TMPPE* | chr3:33134784 | NM_001039770:c.904A>G | p.Asn302Asp | SMP | Yes, not enhanced |
| Proband_132 | Netherlands | Azoospermia | *USH2A* | chr1:216595434 | NM_206933:c.245G>T | p.Arg82Leu | P | Yes, not enhanced |
| Proband_133 | Netherlands | Azoospermia | *EMP1* | chr12:13366446 | NM_001423:c.112G>C | p.Val38Leu | - | Yes, not enhanced |
| Proband_134 | Netherlands | Extreme oligozoospermia | *ERI3* | chr1:44687249 | NM_024066:c.995A>G | p.Gln332Arg | M | Yes, not enhanced |
| Proband_135 | Netherlands | Severe oligozoospermia | *POPDC3* | chr6:105609709 | NM_022361:c.76G>A | p.Glu26Lys | SMP | Yes, enhanced expression in testis |
| Proband_136 | Netherlands | Severe oligoasthenozoospermia | *PLEKHA1* | chr10:124189195 | NM_001001974:c.956C>G | p.Ala319Gly | M | Not expressed |
| | | | *BTAF1* | chr10:93753563 | NM_003972:c.3158T>C | p.Met1053Thr | M | Yes, not enhanced |
| Proband_137 | Netherlands | Azoospermia | *C12orf49* | chr12:117155674 | NM_024738:c.559C>T | p.Arg187Trp | SMP | Yes, not enhanced |
| Proband_138 | Netherlands | Severe oligozoospermia | *GRIP1* | chr12:66849967 | NM_001379345:c.1198C>T | p.Pro400Ser | M | Yes, not enhanced |
| Proband_139 | Netherlands | Extreme oligozoospermia | *RNF223* | chr1:1007489 | NM_001205252:c.458G>A | p.Arg153His | - | Not expressed |
| | | | *ZNF469* | chr16:88494628 | NM_001127464:c.756dup | p.Ala253ArgfsTer118 | N/A | Not expressed |
| | | | *MAP3K3* | chr17:61759150 | NM_203351:c.620C>T | p.Ser207Leu | SMP | Yes, not enhanced |
| | | | *C17orf74* | chr17:7330308 | NM_175734:c.998G>A | p.Arg333Gln | - | Yes, not enhanced |
| | | | *TMPRSS11B* | chr4:69107421 | NM_182502:c.110A>G | p.His37Arg | SP | Not expressed |
| Proband_142 | Netherlands | Azoospermia | *GPR75-ASB3* | chr2:53921057 | NM_016115:c.1333C>T | p.Arg445Cys | SP | Yes, not enhanced |
| Proband_144 | Unknown | Azoospermia | *ODF1* | chr8:103563960 | NM_024410:c.5C>G | p.Ala2Gly | SMP | Yes, enhanced expression in testis |
| Proband_145 | Netherlands | Extreme oligozoospermia | *EXOSC10* | chr1:11136965 | NM_001001998:c.1919dup | p.Asp640GlufsTer2 | N/A | Yes, not enhanced |

| | | | | | | | | |
|---|---|---|---|---|---|---|---|---|
| Proband_146 | Unknown | Severe oligoasthenozoospermia | *OTOA* | chr16:21728238 | NM_144672:c.1499C>T | p.Ala500Val | - | Yes, enhanced expression in testis |
| Proband_148 | Netherlands | Azoospermia | *CRHR1* | chr17:43907477 | NM_001145148:c.452G>A | p.Arg151Gln | SMP | Not expressed |
| | | | *HTT* | chr4:3213834 | NM_002111:c.6595dup | p.Ala2199GlyfsTer9 | N/A | Yes, not enhanced |
| Proband_149 | Netherlands | Azoospermia | *SNED1* | chr2:242012773 | NM_001080437:c.3910A>G | p.Ile1304Val | - | Yes, not enhanced |
| | | | *PCDHB1* | chr5:140431405 | NM_013340:c.354_358dup | p.Glu120GlyfsTer17 | N/A | Yes, not enhanced |
| Proband_150 | Netherlands | Azoospermia | *SPECC1L* | chr22:24761453 | NM_001145468:c.2837G>A | p.Arg946Gln | MP | Yes, not enhanced |
| Proband_153 | Netherlands | Azoospermia | *IQSEC1* | chr3:13008948 | NM_014869:c.4T>A | p.Trp2Arg | MP | Yes, not enhanced |
| Proband_154 | Netherlands | Azoospermia | *ARHGAP33* | chr19:36276183 | NM_052948:c.1814G>A | p.Arg605Gln | SP | Yes, not enhanced |
| | | | *C10orf107* | chr10:63445916 | NM_173554:c.188A>G | p.Asn63Ser | - | Yes, not enhanced |
| Proband_156 | Netherlands | Azoospermia | *LRRN2* | chr1:204587973 | NM_201630:c.1148C>T | p.Thr383Met | SMP | Yes, not enhanced |
| | | | *SRCIN1* | chr17:36714613 | NM_025248:c.2051C>T | p.Ala684Val | M | Not expressed |
| Proband_157 | Netherlands | Azoospermia | *REN* | chr1:204124193 | NM_000537:c.1163C>T | p.Thr388Ile | SM | Not expressed |
| | | | *SIPA1L3* | chr19:38643580 | NM_015073:c.3634G>A | p.Glu1212Lys | SM | Yes, not enhanced |
| Proband_158 | Netherlands | Azoospermia | *TP53TG5* | chr20:44003729 | NM_014477:c.718A>T | p.Thr240Ser | SMP | Yes, enhanced expression in testis |
| | | | *DHX36* | chr3:153994607 | NM_020865:c.2770G>A | p.Asp924Asn | M | Yes, not enhanced |
| | | | *YEATS2* | chr3:183524758 | NM_018023:c.3889C>A | p.Leu1297Ile | M | Yes, not enhanced |
| Proband_160 | Netherlands | Azoospermia | *SDF4* | chr1:1158720 | NM_016176:c.481G>A | p.Glu161Lys | SMP | Yes, not enhanced |
| | | | *ITSN2* | chr2:24522905 | NM_006277:c.1217G>A | p.Arg406Gln | SMP | Not expressed |
| Proband_165 | Netherlands | Extreme oligozoospermia | *MYOF* | chr10:95168662 | NM_013451:c.611G>A | p.Arg204Gln | MP | Yes, not enhanced |
| | | | *RASAL2* | chr1:178435121 | NM_170692:c.3421G>T | p.Glu1141Ter | N/A | Yes, not enhanced |
| Proband_166 | Netherlands | Azoospermia | *C6orf25* | chr6:31691437 | NM_138272:c.83G>A | p.Gly28Glu | SMP | Not expressed |
| Proband_168 | Netherlands | Extreme oligozoospermia | *KLC1* | chr14:104129206 | NM_001130107:c.740del | p.Ser247Ter | N/A | Yes, not enhanced |
| Proband_170 | Netherlands | Azoospermia | *PRPF4B* | chr6:4049340 | NM_003913:c.2026C>T | p.Arg676Cys | SMP | Yes, not enhanced |
| Proband_173 | United Kingdom | Azoospermia | *ZNF629* | chr16:30793127 | NM_001080417:c.2522C>G | p.Pro841Arg | SMP | No expression |
| | | | *CXXC11* | chr2:242812032 | NM_173821:c.124G>A | p.Gly42Ser | - | No expression |
| | | | *IRAK2* | chr3:10255045 | NM_001570:c.683A>T | p.His228Leu | SMP | No expression |
| Proband_178 | United Kingdom | Severe Oligozoospermia | *MICU1* | chr10:74322772 | NM_006077:c.211G>A | p.Gly71Ser | M | Yes, not enhanced |

| | | | | | | | | |
|---|---|---|---|---|---|---|---|---|
| Proband_178 | United Kingdom | Severe Oligozoospermia | *GHRHR* | chr7:31016058 | NM_000823:c.989C>T | p.Ser330Leu | SMP | Not expressed |
| Proband_179 | United Kingdom | Azoospermia | *CELSR2* | chr1:109807146 | NM_001408:c.5360T>C | p.Val1787Ala | SM | Yes, not enhanced |
| | | | *BASP1* | chr5:17275915 | NM_006317:c.590C>T | p.Pro197Leu | M | Not expressed |

* Representation of which tool marks the variant as pathogenic; S=SIFT, M=MUTATIONTASTER, P=POLYPHEN

| Mouse model | Comments | Conclusion variant/gene | Conclusion patient |
|---|---|---|---|
| Yes, Sertoli cell only described[2] | Centrosomal protein regulating centriole engagement and microtubule nucleation (UniProt: Q96SN8). Drosophila models shows meiotic and post-meiotic defects in sperm, mouse model a complete loss of male germ cells. This deficiency occurs in fetal life during the PGC-to-gonocyte transition and seems to result from the dysregulation of mitotic quiescence initiation[2]. In human, CDK5RAP2 is detected in both male and female FGCs[3], which is in line a potential role in maintaining the germ cell pool during embryonic development. Recessive variants described in microcephaly (OMIM: 604804). | Possibly causative | Candidate de novo point mutation (CDK5RAP2) |
| Yes, no infertility described | Catalyzes the hydrolysis of ATP (UniProt: P05023). Involved in sperm capacitation[4]. Gene very intolerant to LoF variation (pLi = 1; LOEUF = 0.12). Missense variants in this gene described in Charcot-Marie-Tooth disease (OMIM: 618036). | Unclear | Multiple candidate genes |
| Yes, no infertility described | Role in actin cytoskeleton for cell migration and adhesion (UniProt: Q9Y4G6). Expressed in spermatogonial stem cells[3] | Unclear | |
| Yes, Sertoli cell only | E3 ubiquitin-protein ligase (UniProt: Q7Z6Z7). Mouse model shows Sertoli cell only phenotype and HUWE1 is required for entry into meiosis and earliest steps in spermatogonial differentiation[5,6]. Interacts with known infertility gene TRIM37[7]. Glutamic acid repeat not well conserved. Expressed in spermatogonial stem cells, differentiating spermatognia and early primary spermatocytes[3]. Missense variants described in intellectual disability (OMIM: 309590) | Possibly causative | |
| Yes, no infertility described | 1 fertile fathers in control cohort shares the exact mutation, Variant predicted to be benign by 3/3 prediction methods. ATP-dependent transporter (UniProt: Q5T3U5). | Not causative | No candidates |
| Yes, no infertility described | Variant predicted to be benign by 3/3 prediction methods. Copper binding function (UniProt: P00450). Interacts with known infertility gene APOA. Known gene for recessive Cerebellar ataxia (OMIM: 604290). | Unlikely causative | |
| Yes, maturation arrest[8] | 4 fertile fathers in control cohort share the exact mutation, Involved in transcription regulation, RNA splicing, RNA transport,DNA repair and damage response (UniProt: P35637). Expressed in spermatogonial stem cells, differentiating spermatognia and early and late primary spermatocytes[3], Known gene for dominant essential tremor (OMIM: 614782) | Not causative | No candidates |
| Yes, no infertility described | Variant predicted to be benign by 2/3 prediction methods. Controls TGF-beta activation by maintaining it in a latent state during storage in extracellular space (UniProt: Q14766). Interacts with known infertility gene APOA. Expressed in spermatogonial stem cells[3] | Unlikely causative | |
| Yes, no infertility described | 1 fertile father in control cohort shares the exact mutation, Outer segement of photoreceptors (UniProt: Q8IWN7). Known gene for dominant occult macular dystrophy (OMIM: 613587) | Not causative | No candidates |
| Yes, no infertility described | Transcriptional regulator recruiting SETDBP1 histone methyltransferase (UniProt: P11308). SETDB1 is crucial for maintainace of embryonic stem cell pools including spermatogonial stem/progenitor cells[9] | Unclear | Candidate de novo point mutation (ERG) |
| Not described | DNA-binding protein involved in cell cycle control (UniProt: Q99459). | Possibly causative | Candidate de novo point mutation (CDC5L) |
| Not described | Actin binding protein with potential role in retina development and exon guidance (UniProt: O14639). Expressed in late primary spermatocytes[10] | Possibly causative | Candidate de novo point mutation (ABLIM1) |
| Not described | 2 fertile fathers in control cohort share the exact same mutation, Variant predicted to be benign by 3/3 prediction methods. Protein function unknown (UniProt: Q96EE4). Expressed in round and elongating spermatids and sperm[10] | Not causative | Candidate de novo point mutation (RASEF) |
| Not described | Binds predominantly GDP, and also GTP (UniProt: Q8IZ41) | Unclear | |
| Yes, female infertility described[11] | Stabilizes microtubules and may regulate actin fiber dynamics through the activation of Rho family GTPases (UniProt: O95996). APC2 is required for oogenesis in mouse and Drosophila[12,13] and is important for asymetric stem cell division of spermatogonial stem cells in Drosophila[14]. Gene known for recessive Sotos syndrome (OMIM: 617169) | Unclear | Candidate de novo point mutation (APC2) |
| Yes, no infertility described | 2 fertile fathers in control cohort share the exact same mutation, Multi-functional cell surface receptor regulating cell adhesion in many diverse developmental processes (UniProt: Q92859) | Not causative | No candidates |
| Yes, no infertility described | Odorant receptor (Potential). May be involved in taste perception (UniProtKB:Q8WZ94) | Unclear | Candidate de novo point mutation (OR5P3) |
| Yes, no infertility described | Physiological suppressor of IKK-epsilon and TBK1 that plays an inhibitory role in virus- and TLR3-triggered IRF3. Inhibits TLR3-mediated activation of interferon-stimulated response elements (ISRE) and the IFN-beta promoter. May act by disrupting the interactions of IKBKE or TBK1 with TICAM1/TRIF, IRF3 and DDX58/RIG-I. Does not inhibit NF-kappa-B activation pathways (UniProtKB:Q9BRV8) | Unclear | Multiple candidate genes |
| Yes, no infertility described | E3 ubiquitin ligase capable of auto-ubiquitination, following phosphorylation by MAP3K3. Potentiates MAP3K3-mediated activation of the NF-kappa-B, JUN/AP1 and DDIT3 transcriptional regulators. Induces apoptosis when overexpressed. Plays a role in the phosphorylation of MAPK1 and/or MAPK3 (UniProtKB: Q6Q0C0) Interacts with known infertility gene TRIM37[7] | Possibly Causative | |
| Not described | Keratin, type I (UniProt: Q14525) | Unlikely causative | Candidate de novo point mutation (ATP8A1) |
| Yes, reduced female infertility | Variant predicted to be benign by 3/3 prediction methods. | Unlikely causative | |
| Yes, no infertility described | Catalytic component of a P4-ATPase flippase complex (UniProt: Q9Y2Q0) | Unclear | |

| | | | |
|---|---|---|---|
| Yes, no infertility described | Variant predicted to be benign by 2/3 prediction methods. May be required for adipogenesis (UniProt: Q8WTT2) | Unlikely causative | No candidates |
| Yes, no infertility described | Regulator of protein export for NES-containing proteins and mRNA nuclear export (UniProt Q9NPJ8). Gene is dispensable for fertility in mice[15]. No LoF variation found in this gene in gnomAD | Unlikely causative | No candidates |
| Yes, no infertility described | Cation channel that gives rise to very low constitutive currents in the absence of activation. The activated channel exhibits selectivity for sodium, and is inhibited by amiloride. (UniProtKB: Q9NY37) | Unclear | Candidate de novo point mutation (ASIC5) |
| Yes, no infertility described | Involved in an inositol phospholipid-based intracellular signaling cascade (UniProt: Q15111) | Unclear | Multiple candidate genes |
| Yes, no infertility described | Acts both as a chaperone in the cytosol and as a chromatin regulator in the nucleus (UniProt: Q99543). Is expressed in PGCs[3]. Gene is required for early embryonic development in mice[16] | Possibly causative | |
| Not described | Variant predicted to be benign by 3/3 prediction methods. Involved in maintaining the homeostasis of cellular nucleotides by catalyzing the interconversion of nucleoside phosphates (UniProt: Q9UIJ7). | Unlikely causative | |
| Yes, reduced female infertility[17] | Cytokine that binds to and signals through the IL1RL1/ST2 receptor which in turn activates NF-kappa-B and MAPK signaling pathways in target cells (PubMed:16286016). Involved in the maturation of Th2 cells inducing the secretion of T-helper type 2-associated cytokines. (UniProtKB: O95760) | Unlikely causative | No candidates |
| Yes, no infertility described | Binds DNA and functions as a transcriptional regulator (UniProtKB: Q9HAZ2). Interacts with known infertility gene CHD7[18]. | Unclear | Multiple candidate genes |
| Yes, maturation arrest[19] | Gene is extremely LoF intolerant (pLi=0.99, LOEUF=0.24) Interacts with known infertility gene CDC14A[20] | Possibly Causative | |
| Yes, infertility of unknown type[21] | Component of the EvC complex that positively regulates ciliary Hedgehog (Hh) signaling (UniProt: P57679). Known gene for recessive Ellis van Creveld syndrome (OMIM: 225500). Gene not intolerant to LoF variation (Pli = 0, LOEUF = 1.06) | Unclear | Candidate de novo LoF mutation (EVC) |
| Yes, no infertility described | Variant predicted to be benign by 2/3 prediction methods. Involved in the regulation of homocysteine metabolism (UniProt: Q93088) | Unlikely causative | |
| Yes, no infertility described | Component of helicase essential for 'once per cell cycle' DNA replication initiation and elongation in eukaryotic cells (UniProt: Q14566). Expressed in PGCs, spermatogonial stem cells and differentiating spermatogonia[3,10]. Gene known for dominant lactase persistence/non-persistence (OMIM: 223100). Knock-down of MCM6 in germ cells in Drosophila resulted in sterility | Possibly causative | Candidate de novo point mutation (MCM6) |
| Not described | Component of a P4-ATPase flippase complex which catalyzes the hydrolysis of ATP (UniProt: Q8TF62) | Unlikely causative | No candidates |
| Not described | Variant predicted to be benign by 2/3 prediction methods. May be involved in transcriptional regulation (UniProt: Q9BSK1). | Unlikely causative | |
| Yes, no infertility described | Variant predicted to be benign by 3/3 prediction methods. Nucleoporin essential for nuclear pore assembly and fusion, nuclear pore spacing, as well as structural integrity (UniProt: Q8TEM1). | Unlikely causative | |
| Not described | Variant predicted to be benign by 2/3 prediction methods. Histone H1 protein binds to linker DNA between nucleosomes forming the macromolecular structure known as the chromatin fiber (UniProt: P16402) | Unlikely causative | No candidates |
| Not described | Fibronectin type III domain-containing protein (UniProt: Q8TC99). Expressed in elongated spermatids and sperm[10] | Unlikely causative | No candidates |
| Not described | Regulates autophagy by playing a role in the reduction of glucose production in an adiponectin- and insulin-dependent manner (UniProt: O94964). Microtubule-associated protein[22]. Very LoF intolerant gene (pLi = 1; LOEUF = 0.19) | Unclear | No candidates |
| Yes, reduced female infertility[23] | Structural component of specialized membrane microdomains known as tetraspanin-enriched microdomains (UniProt: P60033). Gene is important for fertilization[24]. CD82 is expressed in PGCs[3]. Known gene for recessive immunodeficiency (OMIM: 613496) | Possibly causative | Candidate de novo point mutation (CD81) |
| Yes, no infertility described | Phosphoinositide-binding protein which associates with both cell and endoplasmic reticulum (ER) membranes (UniProt: Q9H4L5). Gene not extremely intolerant to LoF variation (pLi = 0; LOEUF = 0.54) | Unclear | Candidate de novo point mutation (OSBPL3) |
| Yes, no infertility described | Variant predicted to be benign by 2/3 prediction methods. ATP-dependent low-affinity peptide transporter which translocates a broad spectrum of peptides from the cytosol to the lysosomal lumen (UniProt: Q9NP78) | Unlikely causative | No candidates |
| Yes, no infertility described | Variant predicted to be benign by 2/3 prediction methods. Receptor for interleukin-12. This subunit is the signaling component coupling to the JAK2/STAT4 pathway (UniProt: Q99665) | Unlikely causative | Candidate de novo point mutation (TOPAZ1) |
| Yes, maturation arrest at the level of spermatocytes[25] | Important for normal spermatogenesis and male fertility. Specifically required for progression to the post-meiotic stages of spermatocyte development (UniProt: Q8N9V7). Gene is abundantly expressed during meiosis[26] | Possibly causative | |
| Not described | 1 fertile father in control cohort shares the exact mutation, Variant predicted to be benign in 2/3 prediction methods, May play a role in innate immunity by inhibiting the antiviral RIG-I signaling pathway (UniProt: Q92503) | Not causative | No candidates |
| Yes, no infertility described | Probable transcription activator for a number of lung-specific genes (UniProt: Q12947). Knock-down of FOXF2 in cysts cells in Drosophila resulted in sterility | Unclear | Candidate de novo point mutation (FOXF2) |
| Yes, no infertility described | Variant predicted to be benign by 2/3 prediction methods. Has E3 ubiquitin ligase activity, promoting ubiquitination and degradation of target proteins (UniProt: O95628) | Unlikely causative | No candidates |
| Not described | Expressed in elongated spermatids[10]. Uncharacterized protein (UniProt Q5SZB4) | Possibly causative | Candidate de novo point mutation (C9orf50) |

| | | | |
|---|---|---|---|
| Yes, no infertility described | Serine/threonine-protein kinase involved in transcription regulation, apoptosis and steroidogenic gene expression (UniProt: Q9H422). Overlaps with previously described CNV[27]. Gene moderately intolerant to LoF variation (pLi = 0.28) | Unclear | Multiple novel candidate genes |
| Not described | Glutamine and serine-rich protein (UniProt Q2KHR3). Overlaps with previously described CNV[27]. Gene very intolerant to LoF variation (pLi = 1) | | |
| Not described | DEP domain-containing protein (UniProt: Q96QD5). Overlaps with previously described CNV[27]. Gene tolerant to LoF variation (pLi = 0) | | |
| Not described | T-complex protein 11-like protein (UniProt: Q9NUJ3). Overlaps with previously described CNV[27]. Gene tolerant to LoF variation (pLi = 0) | | |
| Yes, no infertility described | One of the multiple factors required for polyadenylation and 3'-end cleavage of mammalian pre-mRNAs (UniProt: Q12996). Overlaps with previously described CNV[27]. Gene very intolerant to LoF variation (pLi = 0.98) | | |
| Yes, no infertility described | UPF0606 protein (UniProt: Q6ZVL6). Overlaps with previously described CNV[27]. Gene tolerant to LoF variation (pLi = 0) | | |
| Not described | Tudor domain-containing protein (UniProt: Q5VZ19). Variant predicted to be benign by 2/3 prediction methods. | Unlikely causative | |
| Yes, male infertility of unknown type (MGI:1914535) | As part of the spliceosome, plays a role in pre-mRNA splicing (UniProt: Q6UX04). Known gene for recessive Retinitis pigmentosa (OMIM: 250410) | Possibly causative | |
| Yes, Meiotic arrest[28] | ATPase component of the chromatin remodeling INO80 complex which is involved in transcriptional regulation, DNA replication and DNA repair (UniProtKB: Q9ULG1) Interacts with knocn infertility gene FANCM[29] | Possibly Causative | Candidate de novo point mutation (INO80) |
| Not described | May be involved in transcriptional regulation (UniProt: Q8N972). Gene tolerant to LoF variation (pLi = 0; LOEUF = 1.86) | Unclear | Multiple novel candidate genes |
| Yes, no infertility described | May be responsible for anchoring smooth muscle cells to elastic fibers, and may be involved not only in the formation of the elastic fiber, but also in the processes that regulate vessel assembly (UniProt: Q9Y6C2) | Unclear | |
| Not described | WD repeat-containing protein (UniProt: Q8IZU2) | Unclear | |
| Not described | Variant predicted to be benign by 2/3 prediction methods. May be involved in transcriptional regulation (UniProt: Q5JNZ3) | Unlikely causative | |
| Yes, no infertility described | May play metabolic roles in sperm maturation or fertilization. Phospholipid transfer protein that preferentially selects lipid species containing a palmitoyl or stearoyl chain on the sn-1 and an unsaturated fatty acyl chain (18:1 or 18:2) on the sn-2 position. Able to transfer phosphatidylcholine (PC) and phosphatidyetanolamline (PE) between membranes (UniProtKB: Q9Y365) | Possibly Causative | Multiple novel candidate genes |
| Yes, male infertility of unknown type (MGI:3576497) | Plays a major role in early metanephros and genital development (UniProtKB: Q9C091), Known gene for Renal hypodysplasia (OMIM=617782) Gene is extremely LoF intolerant (pLI=1, LOEUF=0.07) | Possibly Causative | |
| Yes, Meiotic defects[30] | Involved in DNA mismatch repair and meiotic recombination processes. Facilitates crossovers between homologs during meiosis ( UniProtKB: O43196) Gene is not LoF intolerant (pLI=0, LOEUF=0.7) | Unlikely causative | No candidates |
| Not described | Acetolactate synthase-like protein (UniProt A1L0T0) | Unclear | Candidate de novo point mutation (ILVBL) |
| Yes, no infertility described | Sequence-specific transcription factor (By similarity). Regulates multiple developmental processes including brainstem, inner and outer ear, abducens nerve and cardiovascular development and morphogenesis as well as cognition and behavior (UniProt: P49639). | Unclear | Candidate de novo point mutation (HOXA1) |
| Yes, no infertility described | May be involved in transcriptional regulation (UniProtKB: Q86UP3), Gene is extremely LoF intolerant (Pli=1, LOEUF=0.14) | Unclear | Candidate de novo LoF mutation (ZFHX4) |
| Not described | Variant predicted to be benign by 3/3 prediction methods. The B chain of factor XIII is not catalytically active, but is thought to stabilize the A subunits and regulate the rate of transglutaminase formation by thrombin (UniProt: P05160) | Unlikely causative | Candidate de novo point mutation (HNRNPL) |
| Yes, no infertility described | Splicing factor binding to exonic or intronic sites and acting as either an activator or repressor of exon inclusion (UniProt: P14866). | Possibly causative | |
| Yes, no infertility described | Inhibits autoprocessing of CASP1, CASP1-dependent IL1B secretion, PYCARD aggregation and PYCARD-mediated [illegible] | Unlikely causative | No candidates |
| Yes, no infertility described | Component of the PAF1 complex (PAF1C) which has multiple functions during transcription by RNA polymerase II and is implicated in regulation of development and maintenance of embryonic stem cell pluripotency (UniProt Q8WVC0). | Possibly causative | Candidate de novo point mutation (LEO1) |
| Not described | 3 fertile fathers in cohort present with the exact same mutation. Variant predicted to be benign by 2/3 prediction [illegible] | Not causative | No candidates |
| Not described | Targets myosin phosphatase to the actin cytoskeleton. Required for the regulation of the actin cytoskeleton by RhoA and ROCK1 (UniProt: Q6WCQ1) | Unclear | Candidate de novo point mutation (MPRIP) |
| Yes, no infertility described | The heterodimer formed by NGFR and SORCS2 functions as receptor for the precursor forms of NGF (proNGF) and BDNF (proBDNF) (UniProt: Q96PQ0) | Unlikely causative | |
| Yes, no infertility described | Gene is extremely LoF intolerant (Pli=1, LOEUF=0.19) | Unclear | Candidate de novo LoF mutation (TENM2) |
| Yes, no infertility described | Catalyzes the transfer of sulfate to position 4 of the N-acetylgalactosamine (GalNAc) residue of chondroitin and desulfated dermatan sulfate. (UniProtKB: Q9NRB3) | Unlikely causative | No candidates |

| | | | |
|---|---|---|---|
| Yes, female infertility described[32] | Histone demethylase that specifically demethylates both mono- and dimethylated 'Lys-9' of histone H3. May act as a transcription regulator controlling hair biology (via targeting of collagens), neural activity, and cell cycle (UniProt O43593). Known gene for recessive Alopecia universalis (OMIM: 203655) and Atrichia (OMIM: 209500) and dominant Hypotrichosis (OMIM: 146550) | Unlikely causative | Candidate de novo point mutation (SMC2) |
| Yes, no infertility described | Central component of the condensin complex, a complex required for conversion of interphase chromatin into mitotic-like condense chromosomes. The condensin complex probably introduces positive supercoils into relaxed DNA in the presence of type I topoisomerases and converts nicked DNA into positive knotted forms in the presence of type II topoisomerases (UniProt: O95347). Expressed in in spermatogenic stem cells, differentiating spermatogonia and early and late primary spermatocytes[10]. | Possibly causative | |
| Yes, no infertility described | 9 males in gnomAD carry the same variant. Plays a role in the microtubule-dependent coupling of the nucleus and the centrosome. Involved in the processes that regulate centrosome-mediated interkinetic nuclear migration (INM) of neural progenitors (By similarity). May play a role in organizing centrosomal microtubules (UniProt: O95359). Expressed in PGCs[3]. Interacts with known infertility gene AURKC | Unlikely causative | No candidates |
| Yes, no infertility described | A cytochrome P450 monooxygenase involved in the metabolism of endogenous polyunsaturated fatty acids (PUFAs). Mechanistically, uses molecular oxygen inserting one oxygen atom into a substrate, and reducing the second into a water molecule, with two electrons provided by NADPH via cytochrome P450 reductase (UniProtKB: Q9HCS2) | Unlikely causative | Candidate de novo point mutation (RBM5) |
| Yes, spermatid differentiation arrest[33] | Component of the spliceosome A complex. Regulates alternative splicing of a number of mRNAs. May modulate splice site pairing after recruitment of the U1 and U2 snRNPs to the 5' and 3' splice sites of the intron. (UniProtKB: P52756) | Possibly Causative | |
| Yes, no infertility described | Essential for cell viability. TAF9 and TAF9B are involved in transcriptional activation as well as repression of distinct but overlapping sets of genes. (UniProtKB: Q16594) | Unlikely causative | |
| Not described | 8 fertile fathers in cohort present with the exact same mutation, Variant predicted to be benign by 3/3 prediction methods. May act as a downstream effector of CDC42 in cytoskeletal reorganization (UniProt: Q6DT37) | Not causative | Candidate de novo point mutation (RPA1) |
| Yes, no infertility described | As part of the heterotrimeric replication protein A complex (RPA/RP-A), binds and stabilizes single-stranded DNA intermediates, that form during DNA replication or upon DNA stress. It prevents their reannealing and in parallel, recruits and activates different proteins and complexes involved in DNA metabolism (UniProt: P27694). Epxressed in PGCs[3]. Plays an important role in meiotic recombination[34,35]. Interacts with known infertility genes TEX15 and FANCA. | Possibly causative | |
| Yes, no infertility described | Regulator of APC activity during mitotic and meiotic cell cycle, also known as EMI1 (UniProt: Q9UKT4). Required for entry into meiosis and transition from meiosis I to meiosis II in Xenopus oocytes[36]. Expressed in early primary spermatocytes[10]. Gene is extremely intolerant to LoF variation (pLi = 0.97; LOEUF = 0.32). | Possibly causative | Candidate de novo LoF mutation (FBXO5) |
| Yes, no infertility described | Muscle-specific filamin, which plays a central role in muscle cells, probably by functioning as a large actin-cross-linking protein. (UniProtKB: Q14315), Interacts with known infertility gene CFTR[37]. | Unclear | Candidate de novo point mutation (FLNC) |
| Yes, no infertility described | 2 fertile fathers in control cohort share the exact same mutation, Variant predicted to be benign by 2/3 prediction methods. Component of the chromosomal passenger complex (CPC), a complex that acts as a key regulator of mitosis. The CPC complex has essential functions at the centromere in ensuring correct chromosome alignment and segregation and is required for chromatin-induced microtubule stabilization and spindle assembly. Major effector of the TTK kinase in the control of attachment-error-correction and chromosome alignment (UniProt: Q53HL2). Interacts with known infertility gene AURKC | Not causative | No candidates |
| Not described | Variant predicted to be benign by 3/3 prediction methods. Zinc finger CCHC domain-containing protein (UniProt: Q9C0B9) | Unlikely causative | No candidates |
| Yes, no infertility described | AMP deaminase plays a critical role in energy metabolism (UniProtKB: Q01433) | Unclear | Candidate de novo point mutation (AMPD2) |
| Yes, no infertility described | 13 fertile fathers in cohort present with the exact same mutation, This protein specifically binds to the DNA sequence 5'-GGGACTTTCC-3' which is found in the enhancer elements of numerous viral promoters such as those of SV40, CMV, or HIV-1 (UniProtKB: P15822) | Not causative | |
| Yes, abnormal flagellum morphology[38] | Required for correct axoneme development in spermatozoa. Important for normal development of the manchette and sperm head morphology. Essential for male fertility. Plays a role in localization of the intraflagellar transport protein IFT20 to the manchette, suggesting function as an adapter for dynein-mediated protein transport during spermatogenesis (UniProtKB: Q9C093) | Unlikely causative | |
| Not described | 3 fertile fathers in cohort present with the exact same mutation, Variant predicted to be benign by 2/3 prediction methods. Putative adhesion molecule that mediates sialic-acid dependent binding to cells (UniProt: Q96LC7) | Not causative | No candidates |
| Yes, no infertility described | Variant predicted to be benign by 2/3 prediction methods. Seems to act as a glycogen-targeting subunit for PP1. PP1 is essential for cell division, and participates in the regulation of glycogen metabolism, muscle contractility and protein synthesis. Plays an important role in glycogen synthesis but is not essential for insulin activation of glycogen synthase (UniProt: Q16821). Known gene for dominant Insulin resistance (OMIM: 125853) | Unlikely causative | No candidates |

| | | | |
|---|---|---|---|
| Yes, no infertility described | 1 fertile father in control cohort shares the exact mutation,Variant predicted to be benign by 3/3 prediction methods. Sulfotransferase that utilizes 3'-phospho-5'-adenylyl sulfate (PAPS) as sulfonate donor to catalyze the transfer of sulfate to position 6 of non-reducing N-acetylglucosamine (GlcNAc) residues within mucin-associated glycans that ultimately serve as SELL ligands (UniProt: Q8NCG5) | Not causative | Candidate de novo point mutation (STXBP2) |
| Yes, no infertility described | Involved in intracellular vesicle trafficking and vesicle fusion with membranes. Contributes to the granule exocytosis machinery through interaction with soluble N-ethylmaleimide-sensitive factor attachment protein receptor (SNARE) proteins that regulate membrane fusion (UniProt: Q15833). | Unclear | |
| Not described | 3 fertile fathers in cohort present with the exact same mutation, Displays an antiviral effect against flaviviruses such as west Nile virus (WNV) in the presence of OAS1B (UniProt: Q9NUQ8). Expressed in PGCs[3]. | Not causative | No candidates |
| Not described | Transmembrane protein (UniProt: Q0P6H9). Expressed in 10 week old PGCs[3] | Unclear | Candidate de novo point mutation (*U2AF2* ) |
| Not described | Plays a role in pre-mRNA splicing and 3'-end processing. By recruiting PRPF19 and the PRP19C/Prp19 complex/NTC/Nineteen complex to the RNA polymerase II C-terminal domain (CTD), and thereby pre-mRNA, may couple transcription to splicing (UniProt: P26368). Expressed in spermatogonial stem cells, differentiating spermatogonia and early and late primary spermatocytes[3]. Interacts with known infertility gene WT1. Knock-down of U2AF2 in cysts cells in Drosophila resulted in subfertility | Possibly causative | |
| Yes, no infertility described | Helicase that acts as a transcriptional coactivator for a number of nuclear receptors including PPARA, PPARG, THRA, THRB and RXRA (UniProt: Q9BYK8). Interacts with known infertility gene APOA1. | Unclear | Candidate de novo point mutation (HELZ2) |
| Yes, no infertility described | May be a transcriptional repressor of NRL function in photoreceptors (UniProtKB: Q9WTJ4) | Unclear | Candidate de novo point mutation (FIZ1) |
| Yes, no infertility described | Variant predicted to be benign by 3/3 prediction methods. Required for innate immune defense against viruses (UniProtKB: Q7Z434) | Unlikely causative | Candidate de novo point mutation (TMPPE) |
| Not described | Involved in hydrolase activity (UniProtKB - Q6ZT21) | Unclear | |
| Yes, no infertility described | Variant predicted to be benign by 2/3 prediction methods. Involved in hearing and vision as member of the USH2 complex. In the inner ear, required for the maintenance of the hair bundle ankle formation, which connects growing stereocilia in developing cochlear hair cells. In retina photoreceptors, the USH2 complex is required for the maintenance of periciliary membrane complex that seems to play a role in regulating intracellular protein transport (UniProt: O75445). Known gene for recessive Usher syndrome (OMIM: 276901) | Unlikely causative | No candidates |
| Not described | Variant predicted to be benign by 3/3 prediction methods. Epithelial membrane protein (UniProt: P54849) | Unlikely causative | No candidates |
| Yes, no infertility described | Variant predicted to be benign in 2/3 prediction methods. Involved in 3'-5'-exoribonuclease activity (UniProtKB: O43414) | Unlikely causative | No candidates |
| Yes, no infertility described | May play a role in the maintenance of heart function mediated, at least in part, through cAMP-binding (UniProtKB: Q9HBV1) | Unclear | Candidate de novo point mutation (POPDC3) |
| Yes, [39] | Variant predicted to be benign by 2/3 prediction methods, Binds specifically to phosphatidylinositol 3,4-diphosphate (PtdIns3,4P2), but not to other phosphoinositides. May recruit other proteins to the plasma membrane (UniProtKB: Q9HB21) | Unlikely causative | No candidates |
| Yes, no infertility described | Variant predicted to be benign by 2/3 prediction methods, Regulates transcription in association with TATA binding protein (UniProtKB: O14981) | Unlikely causative | |
| Not described | Positively regulates hepatic SREBP signaling pathway by modulating the proper localization of SCAP (SREBP cleavage-activating protein) to the endoplasmic reticulum, thereby controlling the level of functional SCAP (UniProtKB: Q9H741) | Unclear | Candidate de novo point mutation (C12orf49) |
| Yes, no infertility described | 13 males in gnomAD carry the same mutation. Variant predicted to be benign by 2/3 prediction methods, May play a role as a localized scaffold for the assembly of a multiprotein signaling complex and as mediator of the trafficking of its binding partners at specific subcellular location in neurons (UniProtKB: Q9Y3R0) | Unlikely causative | No candidates |
| Yes, no infertility described | 10 males in gnomAD carry the same mutation, Variant predicted to be benign by 3/3 prediction methods, Involved in metal ion binding (UniProtKB: E7ERA6) | Unlikely causative | Multple novel candidate genes |
| Yes, no infertility described | May be involved in transcriptional regulation (UniProtKB: Q96JG9), (Pli score = 0.72, LOEUF= 0.37) | Unclear | |
| Yes, no infertility described | Component of a protein kinase signal transduction cascade. Mediates activation of the NF-kappa-B, AP1 and DDIT3 transcriptional regulators, (UniProtKB: Q99759), Interacts with known infertility gene CDC14A[20] | Possibly Causative | |
| Yes, no infertility described | Variant predicted to be benign by 3/3 prediction methods, Uncharacterized protein (UniProtKB: Q0P670) | Unlikely causative | |
| Yes, no infertility described | Serine protease (UniProtKB: Q86T26) | Unclear | |
| Not described | This protein is involved in the pathway protein ubiquitination, which is part of Protein modification (UniProtKB - A0A6D2WFD3) | Unclear | Candidate de novo point mutation (GPR75-ASB3) |
| Yes, male infertility due to detachment of the sperm head[40,41] | Component of the outer dense fibers (ODF) of spermatozoa. ODF are filamentous structures located on the outside of the axoneme in the midpiece and principal piece of the mammalian sperm tail and may help to maintain the passive elastic structures and elastic recoil of the sperm tail (UniProt: Q14990). ODF1 is reduced in infertile males[42]. ODF1 is expressed in round and elongating spermatids and sperm[10]. | Possibly causative | Candidate de novo point mutation (ODF1) |
| Yes, [43] | Putative catalytic component of the RNA exosome complex which has 3'->5' exoribonuclease activity and participates in a multitude of cellular RNA processing and degradation events. (UniProtKB: Q01780) Gene is not LoF intolerant (Pli=0, LOEUF=0.74) | Unclear | Candidate de novo point mutation (EXOSC10) |

| | | | |
|---|---|---|---|
| Yes, no infertility described | 1 fertile father in control cohort shares the exact mutation, Variant predicted to be benign by 3/3 prediction methods. May act as an adhesion molecule (UniProt: Q7RTW8). Known gene for recessive deafness (OMIM:607039) | Not causative | No candidates |
| Yes, no infertility described | G-protein coupled receptor for CRH (corticotropin-releasing factor) and UCN (urocortin). Has high affinity for CRH and UCN. Ligand binding causes a conformation change that triggers signaling via guanine nucleotide-binding proteins (G proteins) and down-stream effectors, such as adenylate cyclase. Promotes the activation of adenylate cyclase, leading to increased intracellular cAMP levels (UniProtKB: P34998), Interacts with known infertility gene FSHB[44] | Unclear | Multiple novel candidate genes |
| Yes, early stage arrest[45] | May play a role in microtubule-mediated transport or vesicle function.(UniProtKB: P42858) Gene is extremely LoF intolerant (Pli=1, LOEUF=0.18) Gene is associated with Autosomal dominant Huntington disease (OMIM: 613004) | Possibly Causative | |
| Yes, no infertility described | Variant predicted to be benign by 3/3 prediction methods, Involved in calcium ion binding (UniProtKB: Q8TER0) | Unlikely causative | Candidate de novo LoF mutation (PCDHB1) |
| Yes, no infertility described | Potential calcium-dependent cell-adhesion protein. May be involved in the establishment and maintenance of specific neuronal connections in the brain. (UniProtKB: Q9Y5F3), Gene is not LoF intolerant (Pli= 0, LOUEF =1.31) | Unclear | |
| Yes, no infertility described | Involved in cytokinesis and spindle organization. May play a role in actin cytoskeleton organization and microtubule stabilization and hence required for proper cell adhesion and migration. (UniProtKB: Q69YQ0) | Possibly Causative | Candidate de novo point mutation (SPECC1L) |
| Yes, no infertility described | Guanine nucleotide exchange factor for ARF1 and ARF6 (UniProt: Q6DN90). | Unclear | Candidate de novo point mutation (IQSEC1) |
| Yes, no infertility described | May be involved in several stages of intracellular trafficking. (UniProtKB: O14559) | Unclear | Candidate de novo point mutation (ARHGAP33) |
| Yes, male infertility of unknown type (MGI:1920537) | 1 fertile father in control cohort shares the exact mutation, Calcium-binding protein. May be involved in the control of sperm flagellar movement (UniProtKB: Q8IVU9) | Not causative | |
| Yes, no infertility described | Involved in the development and maintenance of excitatory synapse in the vertebrate nervous system. Regulates surface expression of AMPA receptors and instructs the development of functional glutamate release sites (UniProtKB: O43300) | Unclear | Candidate de novo point mutation (LRRN2) |
| Yes, no infertility described | Variant predicted to be benign by 2/3 prediction methods. Acts as a negative regulator of SRC by activating CSK which inhibits SRC activity and downstream signaling, leading to impaired cell spreading and migration (UniProtKB: Q9C0H9) | Unlikely causative | |
| Yes, no infertility described | Renin is a highly specific endopeptidase, whose only known function is to generate angiotensin I from angiotensinogen in the plasma, initiating a cascade of reactions that produce an elevation of blood pressure and increased sodium retention by the kidney (UniProtKB: P00797). Interacts with known infertility genes WT1[46] and CYP21A2[47] | Unclear | Multiple novel candidate genes |
| Yes, no infertility described | Plays a critical role in epithelial cell morphogenesis, polarity, adhesion and cytoskeletal organization in the lens (UniProtKB - O60292) | Unclear | |
| Not described | May play a significant role in p53/TP53-mediating signaling pathway. (UniProtKB: Q9Y2B4) | Possibly Causative | Candidate de novo point mutation (TP53TG5) |
| Yes, no infertility described | Variant predicted to be benign by 2/3 prediction methods, Multifunctional ATP-dependent helicase that unwinds G-quadruplex (G4) structures (UniProtKB: Q9H2U) | Unlikely causative | |
| Yes, no infertility described | Variant predicted to be benign by 2/3 prediction methods, Chromatin reader component of the ATAC complex, a complex with histone acetyltransferase activity on histones H3 and H4 (UniProtKB: Q9ULM3) | Unlikely causative | |
| Yes, no infertility described | May regulate calcium-dependent activities in the endoplasmic reticulum lumen or post-ER compartment (UniProtKB: Q9BRK5) | Possibly Causative | Multiple novel candidate genes |
| Yes, no infertility described | Adapter protein that may provide indirect link between the endocytic membrane traffic and the actin assembly machinery (UniProtKB: Q9NZM3), Interacts with known infertility gene CFTR[37] | Unclear | |
| Yes, no infertility described | 1 fertile father in control cohort shares the exact mutation, Calcium/phospholipid-binding protein that plays a role in the plasmalemma repair mechanism of endothelial cells that permits rapid resealing of membranes disrupted by mechanical stress. Involved in endocytic recycling. (UniProtKB: Q9NZM1) | Not causative | Candidate de novo LoF mutation (RASAL2) |
| Yes, no infertility described | Inhibitory regulator of the Ras-cyclic AMP pathway (UniProtKB: Q9UJF2), Gene relatively intolerant to LoF mutations (pLi score = 0.8, LOEUF=0.34) | Possibly Causative | |
| Yes, no infertility described | Inhibitory receptor that acts as a critical regulator of hematopoietic lineage differentiation, megakaryocyte function and platelet production (UniProtKB: O95866) | Unclear | Candidate de novo point mutation (C6orf25) |
| Yes, no infertility described | Kinesin is a microtubule-associated force-producing protein that may play a role in organelle transport. (UniProtKB: Q07866), Gene is not LoF intolerant (pLi= 0.37, LOUEF =0.41) | Unclear | Candidate de novo LoF mutation (KLC1)) |
| Yes, no infertility described | Has a role in pre-mRNA splicing (UniProtKB: Q13523) | Unclear | Candidate de novo point mutation (PRPF4B) |
| Yes, no infertility described | May be involved in transcriptional regulation. (UniProtKB :Q9UEG4) | Unclear | Candidate de novo point mutation (ZNF629) |
| Yes, no infertility described | Variant predicted to be benign by 3/3 prediction methods. Olfactory receptor binding (UniProtKB: Q14D33) | Unlikely causative | |
| Yes, no infertility described | Binds to the IL-1 type I receptor following IL-1 engagement, triggering intracellular signaling cascades leading to transcriptional up-regulation and mRNA stabilization (UniProtKB: O43187) | Unlikely causative | |
| Yes, no infertility described | Variant predicted to be benign by 2/3 prediction methods. Key regulator of mitochondrial calcium uniporter (MCU) that senses calcium level via its EF-hand domains (UniProtKB : Q9BPX6) | Unlikely causative | No candidates |

| | | | |
|---|---|---|---|
| Yes, infertility of unknown type[1] | Receptor for GRF, coupled to G proteins which activate adenylyl cyclase. Stimulates somatotroph cell growth, growth hormone gene transcription and growth hormone secretion. (UniProtKB: Q02643) | Unlikely causative | No candidates |
| Yes, no infertility described | Receptor that may have an important role in cell/cell signaling during nervous system formation (UniProtKB: Q9HCU4) | Unclear | Candidate de novo point mutation (CELSR2) |
| Yes, no infertility described | Involved in protein domain specific binding (UniProtKB: P80723) | Unlikely causative | |
