## Supplementary Tables 2-5 for "A *de novo* paradigm for male infertility"

**Supplementary table 2: Rare loss-of-function (LoF) mutations observed in additional cohorts of infertile men and fertile control cohorts.**

| Gene | pLI Score | MERGE Cohort of Infertile Men (n=901) | GEMINI Cohort of NOA Men (n=926) | Regeneron Cohort of Infertile Men (n=88) | Italian Cohort of NOA Men (n=48) | Total infertile cohorts (n=1,963) | Fertile Dutch Men (n=5,784) | Fertile Dutch Women (n=5,803) |
|---|---|---|---|---|---|---|---|---|
| ATP1A1 | 1 | N/A | 0 | N/A | 0 | 0 | 0 | 0 |
| CSTF3 | 0.98 | 0 | 0 | 0 | 0 | 0 | 0 | 0 |
| FBXO5 | 0.97 | 0 | 0 | 0 | 0 | 0 | 0 | 0 |
| GREB1L | 1 | 0 | 0 | 0 | 0 | 0 | 0 | 2 |
| HTT | 1 | N/A | 0 | 0 | 0 | 0 | 3 | 3 |
| PPP1R7 | 0.99 | 0 | 0 | 0 | 0 | 0 | 0 | 0 |
| QSER1 | 1 | 0 | 1 | 0 | 0 | 1 | 0 | 1 |
| SOGA1 | 1 | 1 | 0 | 0 | 0 | 1 | 2 | 6 |
| TENM2 | 1 | 0 | 0 | 0 | 0 | 0 | 2 | 2 |
| ZFHX4 | 1 | 0 | 0 | 0 | 0 | 0 | 2 | 2 |

These 10 genes were selected because of a LoF DNM present in the original discovery cohort in these LoF intolerant genes (as defined by a pLI score >0.9). Exome data from four additional cohorts of infertile men as well as control cohorts of fertile men and women were investigated for the presence of LoF mutations in these genes. N/A – Data not available for this gene in WES data.

**Supplementary Table 3: Clinical details of individuals with *RBM5* pathogenic mutations described in this study.**

| Cohort Name | Patient ID | Age of Patient* | Karyotype | Y deletions | Conclusion semen analysis | Conclusion testis histology | Testicular volume left (ml) | Testicular volume right (ml) | Semen conc. (x$10^6$) | Semen volume (ml) | Semen pH | FSH (U/L) | Testicular sperm retrieved | Urological history |
|---|---|---|---|---|---|---|---|---|---|---|---|---|---|---|
| NIJ/NLC Cohort of Patient-Parent Trios | Proband_108 | 40 | 46, XY | None | Severe oligo-zoospermia | No biopsy | NA | NA | 0.5 | 3.2 | 7.5 | 0 | N/A | Unknown |
| NIJ/NLC Cohort of Infertile Men | Proband00282 | 32 | 46, XY | None | Azoospermia | Biopsy compromised | 15 | 15 | 0 | 3.1 | 7.7 | 6.7 | Yes | Cryptorchism with orchidopexy (unknown if unilateral/bilateral) |
| NIJ/NLC Cohort of Infertile Men | Proband00524 | 36 | 46, XY | None | Azoospermia | Hypo-spermatogenesis | 11 | 6 | 0 | 3.4 | 7.5 | 47 | Yes | None |
| MERGE Cohort of Infertile Men | M248 | 43 | 46, XY | None | Azoospermia | Complete SCO | 14 | 27 | 0 | 7.3 | 7.7 | 31.7 | No | None |
| MERGE Cohort of Infertile Men | M2013 | 28 | 46, XY | None | Azoospermia | No biopsy | 7 | 8 | 0 | 3.2 | 8.3 | 27.9 | N/A | None |

* At diagnosis of infertility.

Multiple infertile men from different cohorts were found with a rare pathogenic mutation on *RBM5* in addition to the Proband_108 where a *DNM* on RBM5 was initially identified. Detailed clinical information for the patient from the Geisinger-Regeneron DiscovEHR cohort of infertile men with a rare pathogenic RBM5 mutation is currently not available and was therefore no included in this table.

**Supplementary Table 4: Phasing of *de novo* mutations.**

| | Possibly Causative | Unclear | Unlikely Causative | Not Causative | Total |
|---|---|---|---|---|---|
| Maternal | 1 | 5 | 2 | 2 | 10 |
| Paternal | 6 | 5 | 20 | 12 | 43 |
| Post-Zygotic | 1 | 1 | 3 | 1 | 6 |
| Total | 8 | 11 | 25 | 15 | 59 |

So far, 59 out of 192 variants have undergone phasing to determine the parent of origin. These can then be categorised based on their final classification to investigate the distribution of pathogenic variants with paternal or maternal origin.

**Supplementary table 5:** Long range PCR reaction mixes and running conditions.

**Master mix: BioRad iProof**

Reaction Mix

| | |
|---|---|
| Master mix | 10ul |
| dH2O | 7.4ul |
| DNA | 2ul |
| Primers (combination of equal parts forward and reverse) | 0.6ul |

**Total** 20ul reaction

PCR conditions

| PCR step | Temperature | Time |
|---|---|---|
| **Initial Denaturation** | 98°C | 1 min |
| **Denaturation** | 98°C | 10 sec |
| **Annealing** | 60°C | 15 sec |
| **Extension** | 72°C | 1 min/kb |
| **Final extension** | 72°C | 5 min |

**x30 cycles**

**Master mix: Quantabio RepliQa HiFi ToughMix**

Reaction Mix

| | |
|---|---|
| Master mix | 12.5ul |
| dH2O | 9.0ul |
| DNA | 2.5ul |
| Primers (combination of equal parts forward and reverse) | 1.0ul |

**Total** 25ul reaction

PCR conditions

| PCR step | Temperature | Time |
|---|---|---|
| **Denaturation** | 98°C | 10 sec |
| **Anneal** | 60°C | 1 sec |
| **Extension** | 68°C | ≤ 1 kb: 1 sec 1-10 kb: 5 sec/ kb ≥ 10 kb: 10 sec/kb |

**x30 cycles**

**Master mix: TakaraBio PrimeSTAR GXL**

Reaction Mix

| | |
|---|---|
| PrimeStar GXL DNA Polymerase | 0.5ul |
| dNTP mixture | 2ul |
| 5x PrimeStar GXL Buffer | 5.0ul |
| Primers (combination of equal parts forward and reverse) | 2.0ul |
| DNA | 2.0ul |
| dH2O | 13.5 |

**Total** 25ul reaction

PCR conditions

| PCR step | Temperature | Time |
|---|---|---|
| **Denaturation** | 98°C | 10 sec |
| **Annealing** | 60.0°C | 10 sec |
| **Extension** | 68°C | 1 min/kb |

**x30 cycles**
