## Supplementary Notes 1 for "A *de novo* paradigm for male infertility"

### 1.1 Supplementary notes

Patient recruitment

The patients recruited for this study were enrolled either at the department of Obstetrics and Gynaecology, div, Reproductive Medicine of the Radboud UMC in Nijmegen or at Newcastle clinic.

The department of Obstetrics and Gynecology, div, Reproductive Medicine of the Radboud UMC in Nijmegen is a tertiary referral center for (male) infertility. Approximately 250 couples with a severe male factor are referred annually to our combined fertility and urology consultation for diagnostic evaluation. Prior to this referral, most of the patients have already received urological evaluation by a secondary care center. After referral, all couples are clinically evaluated on the same day by a urologist and gynaecologist or fertility specialist. All males receive routine diagnostic assessment including detailed history and physical examination, scrotal ultrasound, semen analysis and hormone level evaluation of follicle stimulating hormone (FSH), testosterone and often inhibin B (Inh. B). Genetic screening for karyotyping and AZF deletions is performed according to the male subfertility guideline of the Dutch Society of Obstetrics and Gynaecology.

Newcastle Fertility Centre is part of The Newcastle upon Tyne Hospitals NHS Foundation Trust and is a tertiary referral center for patients with infertility. The Andrology Service receives approximately 150 infertility cases per year from secondary and other tertiary referral units across the UK. Male evaluation includes a detailed history and examination, semen analysis and hormone level evaluation of follicle-stimulating hormone (FSH) and testosterone. Genetic screening for karyotyping and AZF deletions is performed according to the male subfertility guideline of the American Urology Association.

Patient selection

A total of 170 patients who presented at the Radboudumc outpatient clinic between July 2007 and September 2018 and a total of 15 patients who presented at the Newcastle Fertility Centre are included in this study.
These patients were all infertile men (< 50 years old), who presented with unexplained (idiopathic) azoospermia or severe oligospermia (sperm concentration of <5 million/ml, by means of 2 semen analyses). Screening results for Y microdeletions, AZF deletions and chromosomal anomalies were negative, and with both parents were living and willing to participate. All patients and their parents gave informed consent prior to participating in the study and undergoing WES.

For the azoospermic patients, care was taken to establish a non-obstructive etiology based on the following parameters: hormone values (FSH, Inh. B), pH and volume of the ejaculate, testicular volume and testicular histology (see Supplementary Notes Tables 1-5)[1]. In some cases these parameters were inconclusive to allow delineation between an obstructive or non-obstructive etiology. Such cases remained included.

Other exclusion criteria were: the occurrence of a language barrier to understand the procedure, unavailability of both parents, a history of chemotherapy or radiotherapy for cancer treatment directly affecting fertility, bilateral orchidectomy, obstructive azoospermia (i.e. previous vasectomy or congenital absence of vas deferens), retrograde ejaculation (an exception was made for a patient who exhibited Sertoli cell only-syndrome) and electro-ejaculation or vibro-ejaculation. Clinical data was obtained from patient records and a paper questionnaire, which was completed by all patients, regarding information on the urological history, medication use and lifestyle.

Semen analysis and definitions

Of all patients, a semen analysis was performed according to WHO 2010 guideline[2]. When spermatozoa were absent after centrifuging the ejaculate, the semen analysis was repeated at a later date. Reference values and nomenclature were used according to the WHO guidelines[2] except for oligozoospermia which was sub-classified according to concentrations levels:

- Severe oligozoospermia: 0.1 - 5 million sperm/ml

- Extreme oligozoospermia: < 0.1 million sperm/ml

Motility and morphology parameters of spermatozoa were not determined when their concentration was below 0.1 million sperm/ml.

Cohort description

A total of 185 infertile men were included. Clinical characteristics of all patients are provided in Supplementary Table 1. The average age of included men was 33 years (range 20 to 49 years). Most men were of Dutch or British descent (n=163 and n=15 respectively). One male was from the United Arab Emirates, one male was from Saint Martin (Caribbean Islands) and of four men their descent was unknown. In 52 men (28%), there was a family history of genetic abnormalities or infertility. More than half of the patients (101/179, 56%, 6 unknown) had an uneventful urological history. Most reported abnormalities in urological history were cryptorchidism (52/78, 67%) and/or inguinal hernia (17/78, 22%) and/or the occurrence of a varicocele or hydrocele (14/78, 18%). Scrotal ultrasound was abnormal in 58% of patients which this data was available (77/132) most often due to varicocele or hydrocele (40/77, 52%), and/or cysts in the epididymis or testis (21/77, 27%) and/or (micro)calcifications (16/77, 21%). One patient had an undefinable tumour in the left testis, which turned out to be benign.

Semen analysis diagnosed azoospermia in 111 of 185 men (60%) and 74 men with oligozoospermia (40%) (Supplementary Notes Table 1). The oligozoospermic men were then classified as extreme oligozoospermia in 41 cases (55%) and severe oligozoospermia in 33 (45%). When sperm mobility and morphology was taken into account for the men with severe oligozoospermia (since these parameters cannot be reliably be determined in case of extreme oligozoospermia) the following distribution was seen: 13 men presented with oligoasthenoteratozoospermia, 10 with oligoasthenozoospermia, 5 with oligoteratozoospermia and 5 with oligozoospermia.

In total, 6 PESE procedures and 118 TESE procedures were performed. Sperm was found in 65 (55%) of the TESE procedures, in 59 of these couples an ICSI-TESE was performed, which led to a pregnancy in 43 of these treatments (73%, ultrasound at 7-8 weeks). Five out of six ICSI-PESE couples achieved a pregnancy. In addition, 51 of the 74 patients with severe to extreme oligozoospermia had an ICSI with ejaculated sperm. A total of 44 (86%) of these couples achieved a pregnancy. Overall, 92 out of 185 couples (49%) of which the male presented with severe infertility achieved a pregnancy during their treatment. All the medical details and treatment outcomes are provided in Supplementary Notes Table 2. A flowchart of diagnostic and clinical outcomes is presented in Supplementary Figure 1.

Processing of the TESE biopsies and evaluation of spermatogenesis

TESE surgery and subsequent biopsy processing was performed as described in Hessel et al 2013[3]. An overview is shown in Supplementary notes Figure 1. Briefly, together with the testicular biopsy for sperm retrieval a biopsy for diagnostic purposes was taken. The latter was fixed overnight in formaldehyde and paraffin embedded. For histological evaluation of spermatogenesis H&E stainings were made. Presence of a malignancy was determined by immunohistochemistry with OCT3/4- and PLAP-antibodies.

Spermatozoa were isolated from the testicular biopsy by repetitive mincing of the biopsy with two sterilized microscope slides. The obtained testicular cell suspension was inspected by two laboratory technicians. When spermatozoa were observed, the samples were cryo-preserved. At the Radboudumc a droplet from the testicular cell suspension was placed on a slide, let to dry and eventually stained with Giemsa for cytological analysis[4]. The diagnostic biopsy was embedded in a paraffin block. For the samples from the Radboudumc the remaining scrapes of the biopsy was added to the diagnostic biopsy[4].

Description of spermatogenesis

The evaluation of spermatogenesis was based on the histological evaluation of H&E stained sections and the clinical result of the TESE (during which the processed biopsy is inspected for presence of spermatozoa and the material is cryopreserved or not). For the samples collected at the Radboudumc cytological insight was obtained from Giemsa-stained slides of the cell suspension generated from the testicular biopsy. The histological and cytological evaluations were performed on coded samples. Eventually the conclusions of the sub-evaluations (histology, cytology and clinical result TESE) were combined with the outcomes of the patient's semen analysis. Based on all these aspects, a description and conclusion of spermatogenesis for each patient was generated (see Supplementary Notes Table 4).

Completeness of sampling

In 84 out 118 patients that underwent TESE, both histology and cytology samples were available (71%), for 31 patients only histology (n=12, 10%) or cytology (n=19, 16%) was available and could be studied. In three cases (2%) both were unavailable and only the observation whether spermatozoa were detected in the testicular cell suspension remained as an indication for the functionality of spermatogenesis in these patients.

Histological evaluation

In each biopsy a minimum of 100 seminiferous tubules were inspected by two researchers for the presence of germ cells, Sertoli cells and possible abnormalities in the tissue like hyalinization[5]. The flowchart below (Supplementary Notes Figure 2) schematically represents the categorization which was used to diagnose the biopsies. The functionality of spermatogenesis was denoted in six main classifications and six sub-classifications (see Supplementary Notes Table 3). The classification framework largely followed the one described in McLachlan et al[5]. An additional classification 'SCO/minimal-Germ Cell Arrest' was added describing biopsies that predominantly (>95%) contain tubules with only Sertoli cells but also contain an occasional tubule that show germ cells (though no elongating spermatids). If elongating spermatids were present in such a biopsy, it was classified as hypospermatogenesis (HS). The introduction of this classification allows for the distinction between samples showing germ cell arrest with overall absence of germ cells from the seminiferous epithelium and samples with a germ cell arrest without major loss of germ cells from the epithelium.

When germ cells showed a more or less uniform arrest in the seminiferous epithelium the sample was categorized as Germ Cell Arrest (GCA). A subclassification was made according to the stages of spermatogenesis: spermatogonial proliferation, meiosis and spermiogenesis. The stage which is predominant is the denominator

The term hypospermatogenesis (HS) was used for biopsies in which germ cells density have decreased in the seminiferous epithelium but elongating spermatids are present. A reduced number of germ cells in the epithelium can stem from different scenarios: germ cell progenitors might have arrived in the fetal testis in reduced numbers, unable to populate the entire testis, spermatogonia might have been depleted during life or a loss of maturing germ cells is seen during spermatogenesis. The first two scenarios will lead to seminiferous tubules where only Sertoli cells are present. The latter scenario will result in epithelium with ongoing spermatogenesis in which germ cell cohorts are lost. Samples resembling this scenario were categorized as 'Hypospermatogenesis with partial germ cell arrest' (HS-pGCA). Since the degree of germ cell loss varies between samples we introduced 'mild spermatogenesis' (HS-mild) and 'severe hypospermatogenesis' (HS-severe). Mild hypospermatogenesis was defined as the presence of elongating spermatids in 10-50% of tubules. Severe hypospermatogenesis was defined as the presence of elongating spermatids in less than 10% of the tubules. When more than 50% of the tubules did show elongating spermatids, the biopsy was classified as 'complete spermatogenesis' (CS). The 50% cut-off was based on an analysis of testicular biopsies from men that previously had undergone a vasectomy (unpublished data F. Tüttelmann). It should be stressed that complete spermatogenesis does not imply that spermatogenesis was normal, such samples could still show overall reduction in germ cells in the epithelium. Hyalinization was reported if it exceeded >50% of the tubules. Supplementary Notes Table 4 contains a description of all TESE biopsies.

Cytological evaluation

In an air-dried, Giemsa-stained preparation of the TESE-biopsy derived cell suspension, spermatocytes, spermatozoa and Sertoli cells can be readily identified[4]. Since this biopsy is from a different testicular territory as the biopsy used for histological sections it potentially yields additional information on the status of spermatogenesis in the patient. In addition, the ratio of spermatocytes over spermatozoa and the ratio of spermatozoa over Sertoli cells can be used to gain extra information on efficiency of germ cell maturation and sperm production respectively[4,6].

Each slide was scanned by two researchers (GvdH and LR) for presence of spermatocytes, spermatozoa and Sertoli cells. A minimal number of 400 cells were counted in randomly chosen microscopic fields. From these cell counts aforementioned ratios were determined and samples classified. A spermatocytes/spermatozoa ratio > 0.70 was indicative for a partial germ cell arrest. A spermatozoa/ Sertoli cell ratio <28 was indicative for hypospermatogenesis[4].

In some samples both ratios were shifted, indicating that germ cells present had problems to fully finalize spermatogenesis. In the cytological analysis such samples were labelled 'Hypospermatogenesis with partial-Germ Cell Arrest'. Combined with histological information the characterisation of the sample was then reduced to either hypospermatogenesis or partial Germ Cell arrest.

When the combined number of spermatocytes and spermatozoa was below 5% of the total cell count, its ratio was regarded as uninformative and the sample was categorized as 'Hypospermatogenesis'. All results are available in Supplementary Notes Table 5.

Combining observations

In most samples the histology, cytology and clinical TESE result were congruent. Whenever these observations were incongruent, the least severe diagnosis was given. For example, see Proband_122: based on both histology and cytology preparations spermatogenesis was arrested at the spermatocytes stage, indicating a germ cell arrest. However, the clinical TESE result implied the presence of spermatozoa, therefore the final description for the testicular biopsy was 'Hypospermatogenesis with partial Germ Cell Arrest'.


1 Tournaye, H., Krausz, C. & Oates, R. D. Concepts in diagnosis and therapy for male reproductive impairment. *The lancet. Diabetes & endocrinology* **5**, 554-564, doi:10.1016/s2213-8587(16)30043-2 (2017).

2 World Health Organization. WHO laboratory manual for the examination and processing of human semen. (2010).

3 Hessel, M. *et al.* A novel cell-processing method 'AgarCytos' in conjunction with OCT3/4 and PLAP to detect intratubular germ cell neoplasia in non-obstructive azoospermia using remnants of testicular sperm extraction specimens. *Human reproduction (Oxford, England)* **28**, 2608-2620, doi:10.1093/humrep/det311 (2013).

4 Hessel, M. *et al.* Cytological evaluation of spermatogenesis: a novel and simple diagnostic method to assess spermatogenesis in non-obstructive azoospermia using testicular sperm extraction specimens. *Andrology* **3**, 481-490, doi:10.1111/andr.12023 (2015).

5 McLachlan, R. I., Rajpert-De Meyts, E., Hoei-Hansen, C. E., de Kretser, D. M. & Skakkebaek, N. E. Histological evaluation of the human testis--approaches to optimizing the clinical value of the assessment: mini review. *Human reproduction (Oxford, England)* **22**, 2-16, doi:10.1093/humrep/del279 (2007).

6 Skakkebaek, N. E. & Heller, C. G. Quantification of human seminiferous epithelium. I. Histological studies in twenty-one fertile men with normal chromosome complements. *J Reprod Fertil* **32**, 379-389, doi:10.1530/jrf.0.0320379 (1973).

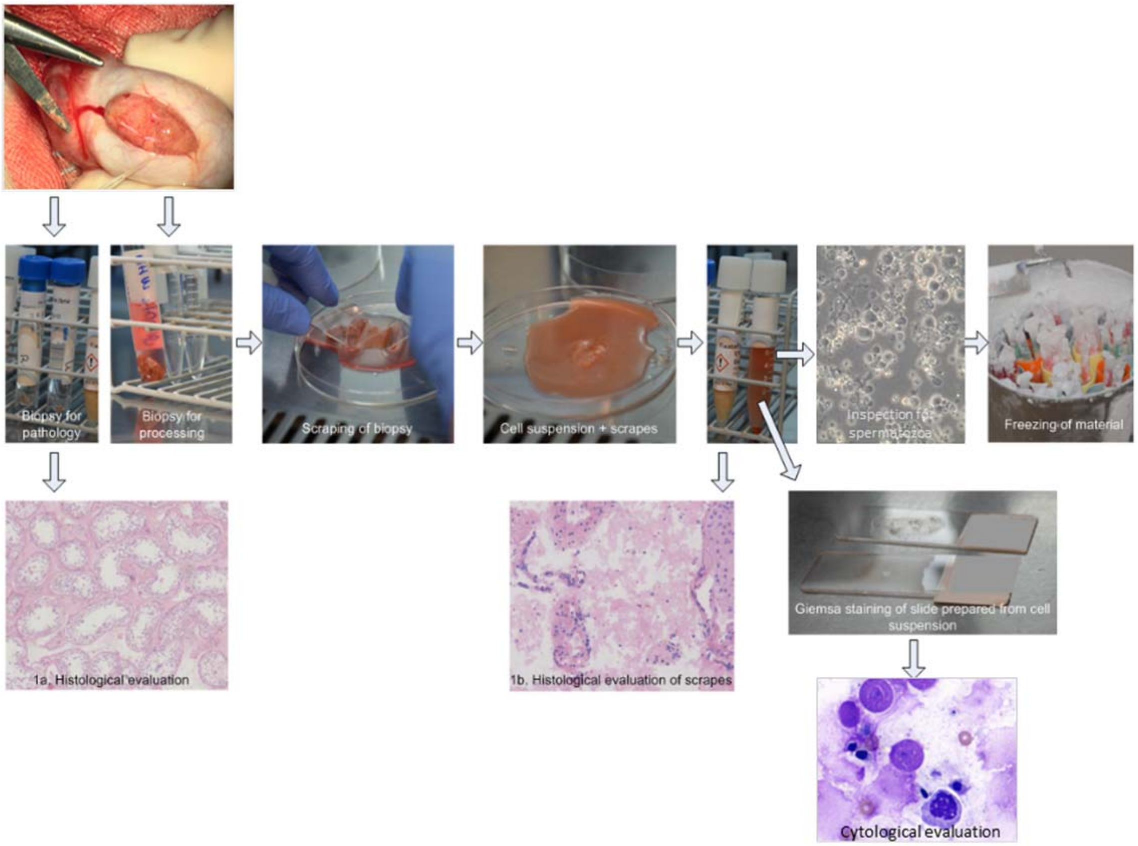


**Supplementary Notes Figure 1: Overview of laboratory processing of testicular biopsies and evaluation of spermatogenesis.** Evaluation of spermatogenesis as presented in this study is based on two different sources: 1. Histological slides made from an unprocessed biopsy (1a) and/or from TESE scrapes (1b) and 2. Outcome of inspection by laboratory technician of sample from cell suspension derived from testicular biopsy. For samples from the Radboudumc a cytological evaluation was available for which GIEMSA stained slides with cells derived from the processed testicular biopsy were prepared.

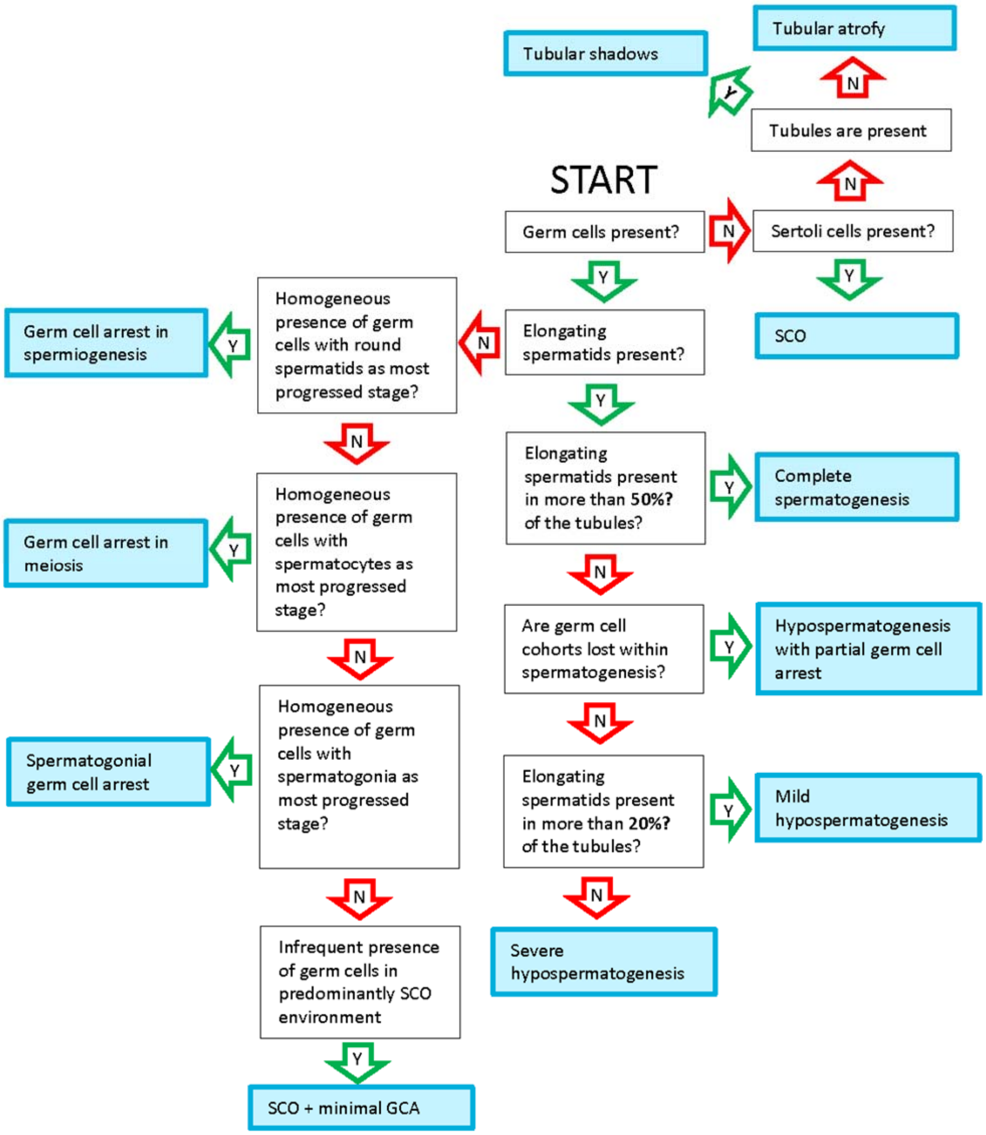


**Supplementary Figure 2: Flowchart for classification of testis histology**
