## Supplementary Notes 2 for "A *de novo* paradigm for male infertility"

### Supplemental information on the authorship: Genetics of Male Infertility Initiative (GEMINI) consortium and its contributors

#### GEMINI participating centers:

Department of Genetics, Oregon National Primate Research Center, Oregon Health & Science University, Beaverton, OR, USA
**Donald F. Conrad, Liina Nagirnaja**

Andrology and IVF Laboratory, Department of Surgery (Urology), University of Utah School of Medicine, Salt Lake City, UT, USA
**Kenneth I. Aston**, **Douglas T. Carrell**, **James M. Hotaling, Timothy G. Jenkins**

1) Hudson Institute of Medical Research and the Department of Obstetrics and Gynaecology, Monash University, Clayton, Victoria, Australia; 2) Monash IVF and the Hudson Institute of Medical Research, Clayton, Victoria, Australia
**Rob McLachlan**

School of Biological Sciences, Monash University, Clayton, Victoria, Australia
**Moira K. O'Bryan**

Department of Urology, Weill Cornell Medicine, New York, NY, USA
**Peter N. Schlegel**

Department of Urology, Stanford University School of Medicine, Stanford, CA 94305, USA
**Michael L. Eisenberg**

Department of Urology, Medical College of Wisconsin, Milwaukee, WI, 53226, USA
**Jay I. Sandlow**

Washington University in St Louis, School of Medicine, St Louis, MO, USA
**Emily S. Jungheim**, **Kenan R. Omurtag**

1) i3S - Instituto de Investigação e Inovação em Saúde , Universidade do University of Porto
2) IPATIMUP - Instituto de Patologia e Imunologia Molecular da Universidade do Porto, Porto, Portugal
3) Serviço de Genética, Departamento de Patologia, Faculdade de Medicina da Universidade do Porto, Porto, Portugal
**Alexandra M. Lopes[1,2], Susana Seixas[1,2], Filipa Carvalho[1,3], Susana Fernandes[1,3], Alberto Barros[1,3]**

1) Departamento de Genética Humana, Instituto Nacional de Saúde Dr Ricardo Jorge, Lisboa, Portugal

2) ToxOmics, Faculdade de Ciências Médicas, Universidade Nova de Lisboa, Portugal
3) Centro de Medicina Reprodutiva, Maternidade Dr. Alfredo da Costa, Lisboa, Portugal
**João Gonçalves[1,2], Iris Caetano1, Graça Pinto[3], Sónia Correia[3]**

Institute of Biomedicine and Translational Medicine, University of Tartu, 51010 Tartu, Estonia
**Maris Laan**

Andrology Center, Tartu University Hospital, 50406 Tartu, Estonia
**Margus Punab**

Department of Growth and Reproduction, Rigshospitalet, University of Copenhagen, Copenhagen, Denmark
**Ewa Rajpert-De Meyts, Niels Jørgensen, Kristian Almstrup**

1) Department of Experimental and Clinical Biomedical Sciences, University of Florence, Florence, Italy; 2) Andrology Department, Fundacio Puigvert, Instituto de Investigaciones Biomédicas Sant Pau (IIB-Sant Pau), Barcelona, Spain
**Csilla G. Krausz**

Division of Urology, Department of Surgery, Mount Sinai Hospital, University of Toronto, Toronto, ON, Canada
**Keith A. Jarvi**
